# Ultra-deep Sequencing of Hadza Hunter-Gatherers Recovers Vanishing Gut Microbes

**DOI:** 10.1101/2022.03.30.486478

**Authors:** Bryan D. Merrill, Matthew M. Carter, Matthew R. Olm, Dylan Dahan, Surya Tripathi, Sean P. Spencer, Brian Yu, Sunit Jain, Norma Neff, Aashish R. Jha, Erica D. Sonnenburg, Justin L. Sonnenburg

**Affiliations:** Department of Microbiology and Immunology, Stanford University School of Medicine, Stanford, CA, USA; Chan Zuckerberg Biohub, San Francisco, CA, USA; Genetic Heritage Group, Program in Biology, New York University Abu Dhabi, Abu Dhabi, UAE; Center for Human Microbiome Studies, Stanford University School of Medicine, Stanford, CA, USA

## Abstract

The gut microbiome is a key modulator of immune and metabolic health. Human microbiome data is biased towards industrialized populations, providing limited understanding of the distinct and diverse non-industrialized microbiomes. Here, we performed ultra-deep metagenomic sequencing and strain cultivation on 351 fecal samples from the Hadza, hunter-gatherers in Tanzania, and comparative populations in Nepal and California. We recover 94,971 total genomes of bacteria, archaea, bacteriophages, and eukaryotes, 43% of which are absent from existing unified datasets. Analysis of *in situ* growth rates, genetic *pN/pS* signatures, high-resolution strain tracking, and 124 gut-resident species vanishing in industrialized populations reveals differentiating dynamics of the Hadza gut microbiome. Industrialized gut microbes are enriched in genes associated with oxidative stress, possibly a result of microbiome adaptation to inflammatory processes. This unparalleled view of the Hadza gut microbiome provides a valuable resource that expands our understanding of microbes capable of colonizing the human gut and clarifies the extensive perturbation brought on by the industrialized lifestyle.

## Introduction

The gut microbiome is increasingly recognized as a critical aspect of human health. While microbiome composition varies profoundly across global lifestyles, microbiome studies are heavily biased towards western industrialized populations (Abdill et al., 2022). Industrialized populations are characterized by low microbiome diversity, and aspects of lifestyle, including i) consumption of highly-processed foods, ii) high rates of antibiotic administration, iii) birth via cesarean section, iv) sanitation of the living environment, and v) reduced physical contact with animals and soil have been hypothesized to mediate this reduced diversity (Sonnenburg and Sonnenburg, 2019b). These aspects are absent from the lifestyle of non-industrialized human populations, including hunter-gatherers who harbor extremely high microbiome diversity (Smits et al., 2017). The transition to an industrialized microbiome is observed in immigrants to the United States of America, supporting a causal role of lifestyle (Vangay et al., 2018).

Groups of microbial taxa that are specifically associated with industrialized and non-industrialized populations are referred to as BloSSUM (Bloom or Selected in Societies of Urbanization/Modernization) and VANISH (Volatile and/or Associated Negatively with Industrialized Societies of Humans) taxa, respectively (Blaser, 2017; Clemente et al., 2015; Martínez et al., 2015; Modi et al., 2014; Mueller et al., 2015; Sonnenburg and Sonnenburg, 2019a; Yatsunenko et al., 2012). Analysis of coprolites supports the view that ancient microbiomes more closely resemble the modern non-industrialized microbiome than the industrialized microbiome (Wibowo et al., 2021). Human-associated microbial lineages have been passed across hominid generations over evolutionary time (Linz et al., 2007; Moeller et al., 2014), raising the possibility that human biology has become reliant upon functions and cues that these VANISH microbes provide (Blaser and Falkow, 2009).

Our current understanding of the VANISH taxa is crude and primarily based on 16S rRNA sequencing (Sonnenburg and Sonnenburg, 2019b), and therefore lacks phylogenetic resolution and genomic/functional insight. A higher-resolution view, including an understanding of VANISH functional capacity, growth dynamics, and dispersal patterns, is needed to understand microbiome change induced by the industrialized lifestyle. Further, recent efforts to establish comprehensive databases of gut-associated genomes and genes have shown that populations living non-industrial lifestyles still have not been sequenced sufficiently to capture the extent of microbiome novelty (Almeida et al., 2019, 2020; Nayfach et al., 2019; Olm et al., 2020; Pasolli et al., 2019).

Metagenomic sequencing has transformed our ability to understand microbes without cultivation. Most modern human microbiome studies use relatively shallow sequencing, but deep sequencing improves *de novo* genome recovery (including from microbial eukaryotes (Olm et al., 2019a)) and allows the use of recently-developed techniques such as *in situ* growth rate prediction and high-precision strain-tracking and microdiversity analysis (Brown et al., 2016; Olm et al., 2021). Further, more complex microbiomes require deeper sequencing for tasks such as metagenomic assembly. Therefore, in addition to reconciling the decreased representation of non-industrialized lifestyle populations in general (Abdill et al., 2022), there is a key need for deep metagenomic sequencing from these populations to better understand their complex and diverse microbiomes.

Here we present ultra-deep metagenomic sequencing and high-resolution analysis of the Hadza hunter-gatherer gut microbiome. We report 9.4 Tbp of metagenomic sequencing data generated from 351 Hadza fecal samples, high-quality metagenomic assemblies, 83,044 *de novo* metagenomic assembled genomes (MAGs) from 4 domains of life, and the results from numerous state-of-the-art bioinformatic techniques. Among these metagenomes is the most deeply-sequenced human gut metagenome to date (210 Gbp). Crucially, because the sequencing depth generated for these Hadza samples is so much higher than previous studies, we also performed deep sequencing on Nepali and Californian populations to enable microbiome lifestyle comparisons without the need for sequence rarefaction. The data generated allow us to make several key insights into the Hadza gut microbiome and the impacts of industrialization. All data generated and the results of all analyses performed are made freely publicly available as a resource for future study by other scientists, including i) metagenomic sequencing reads, assemblies, and MAGs, ii) prevalence data for 5,755 species-level representative genomes across 22 global microbiome studies, iii) isolate genome reads, genomes, and isolate-to-MAG comparisons, and iv) organized metadata.

## Results

### Generating a Vast Resource of Hadza Gut Microbiome Data

The Hadza reside near Lake Eyasi in the central Rift Valley of Tanzania. They live in bush camps of approximately 5 to 30 people, move between camps approximately every 4 months, primarily drink from water springs and streams, and eat a diet that includes foraged tubers, berries, honey, and hunted animals (Marlowe, 2010). They are among the last remaining populations in Africa that continue a form of the ancestral foraging legacy of our human species.

We performed metagenomic sequencing on stool samples collected from 167 Hadza individuals (including 33 infants and 6 mothers (Olm et al., 2022)) between September 2013 and August 2014 (Fragiadakis et al., 2019; Smits et al., 2017) **(Fig. 1A; Supplementary Table 1**). Of these, 101 individuals were sampled once and 66 individuals were sampled longitudinally **(Fig. 1A)**. DNA extraction was performed using the MoBio PowerSoil kit (n=318), phenol chloroform extraction (n=38), or both (n=32), and the resulting DNA was subjected to paired-end Illumina shotgun sequencing **(Fig. 1B)**. Extraction methods did not have a statistically significant effect on the number of genomes detected per sample (**Fig. S1**).

**Fig. 1.**
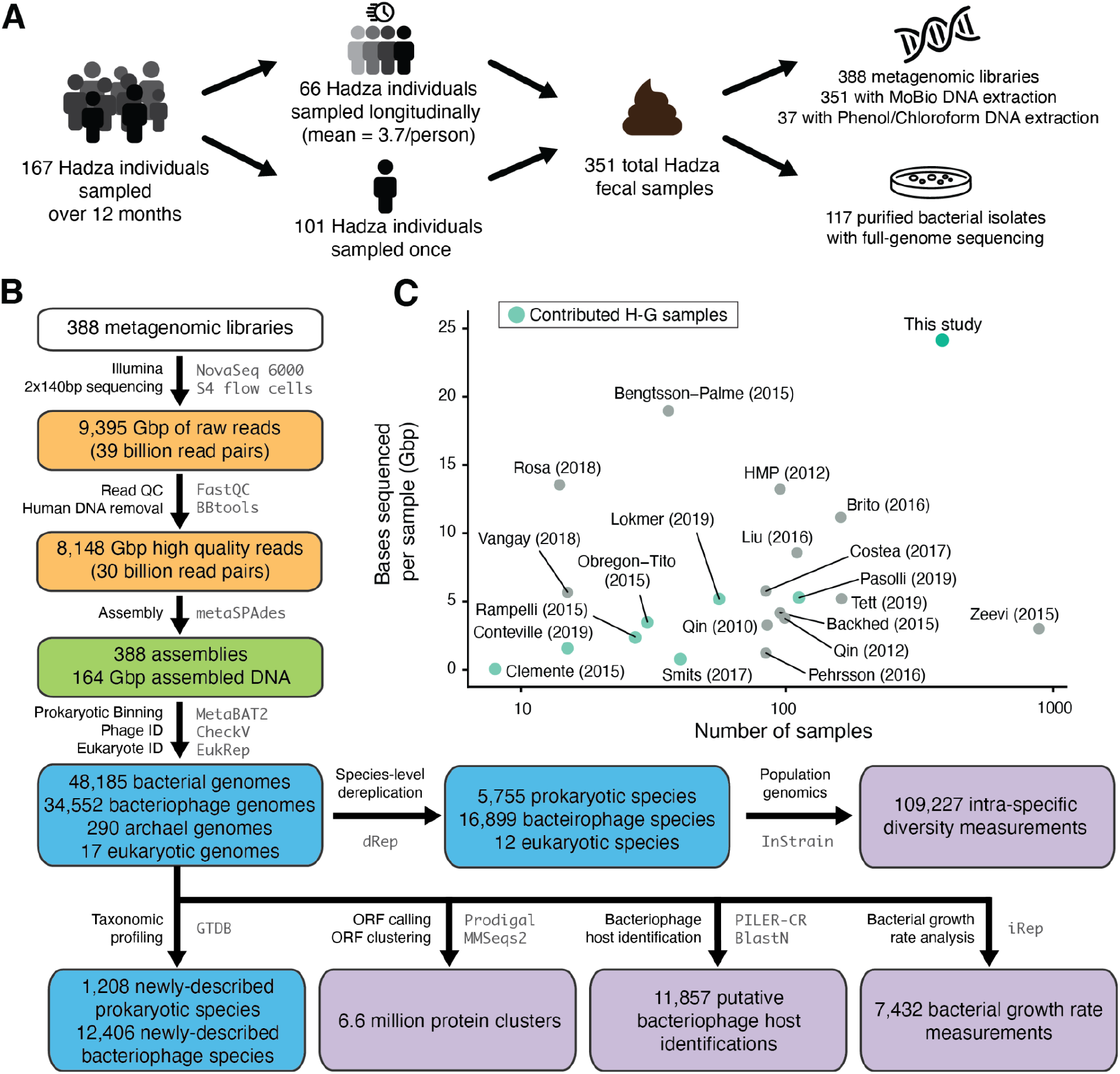
A vast resource of Hadza gut microbiome data. **(A)** Overview of sample collection for shotgun metagenomic sequencing of Hadza fecal samples. **(B)** Summary of the computational workflow, tools used, and primary data generated from Hadza stool samples. **(C)** Number of samples versus the number of bases sequenced per sample for 21 previously published human gut metagenomic data sets and the present study.

A total of 9,395 giga base-pairs (Gbp) of metagenomic data were generated from these 388 Hadza metagenomes (range = 0.7 – 210.3 Gbp, avg = 21.0 Gbp, std dev = 14.5 Gbp). Both the sequencing depth per sample and the overall number of samples sequenced in this study are exceptional relative to previously reported human microbiome metagenomic studies **(Fig. 1C)**. Using multi-domain assembly, binning, and read-mapping **(Fig. 1B)**, we recovered 48,475 bacterial and archaea MAGs (≥ 50% completeness and ≤ 10% contamination; medium quality or better by MiMAG standards (Bowers et al., 2017)), 17 eukaryote MAGs (≥ 50% completeness and ≤ 15% contamination according to EukCC (Saary et al., 2020)), and 4,552 bacteriophage MAGs (medium-quality or better according to CheckV (Nayfach et al., 2021a)) (**Table S2**). We performed numerous diversity, taxonomic, and functional analyses on these recovered genomes, as outlined in **Fig. 1B** and described below.

Many genomes recovered from the Hadza come from species that are absent from the Unified Human Gastrointestinal Genome (UHGG) database (Almeida et al., 2020), the Genome Taxonomy Database (GTDB) (Parks et al., 2022) (**Fig. 2A**), and the Metagenomic Gut Virus (MGV) catalog (Nayfach et al., 2021b) (**Fig. 2B**). MAGs recovered from the Hadza expand the Unified Human Gastrointestinal Genome (UHGG, v1) database (Almeida et al., 2020) bacterial and archaeal species count by 25.4% and 14.3%, respectively, and the Metagenomic Gut Virus (MGV) catalog (Nayfach et al., 2021b) viral species count by 23.7%. The majority of eukaryotic genomes recovered from the Hadza are from the genus *Blastocystis* (n=10), a prevalent member of the mammalian gut microbiota (Clark et al., 2013). Of the 7 other eukaryotic genomes recovered from the Hadza gut, one is a remarkably large and complete genome of a stingless bee (232 megabase pairs and 92.3% complete), the honey and larvae of which are known to be consumed by the Hadza (Marlowe et al., 2014), and four are novel *Amoebae* (n=2) and *Trepomonas* (n=2) genomes **(Fig. 2C)**. While a comprehensive genome database does not yet exist for eukaryotes known to colonize the human gut, genomes from these species are not present in NCBI GenBank (a repository of genomes sequenced from all environments) (NCBI Resource Coordinators, 2017). Finally, over half (59.7%) of the 6.6 million protein families (clustered at 95% amino acid identity) found in Hadza gut microbes are absent from the UHGP-95 protein database (Almeida et al., 2020), a collection of all proteins from genomes in UHGG **(Fig. 2D)**. Together these data highlight the exceptional species- and gene-level novelty elucidated by deep sequencing within this single study.

**Fig. 2.**
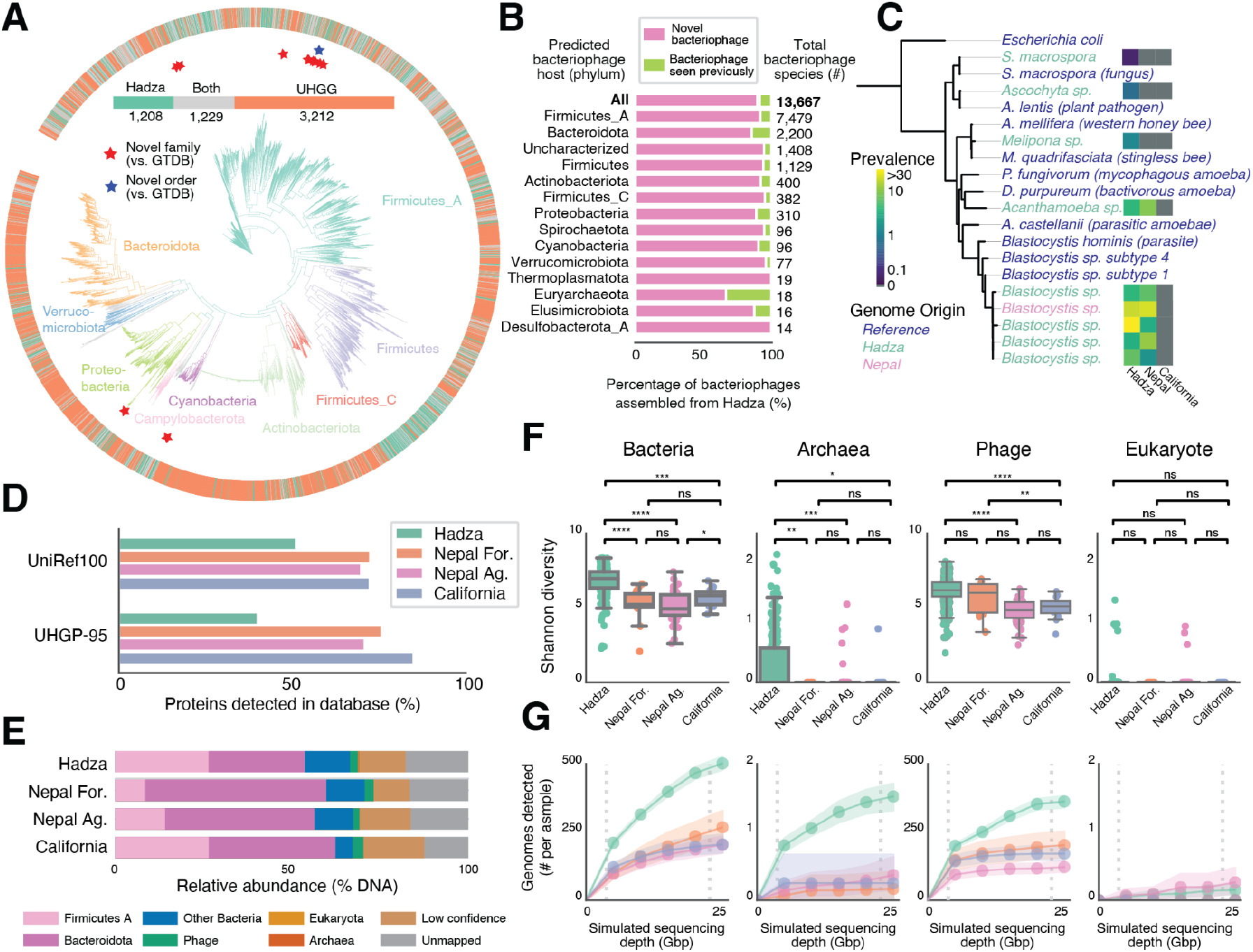
The Hadza gut microbiota contains substantial multi-domain novelty. **(A)** Phylogenetic tree of bacterial species-level representative genomes (SRGs) from Hadza and UHGG based on bacterial single copy gene alignment; branch colors correspond to phyla. SRGs from species-level groups consisting of only genomes assembled from the Hadza or only UHGG are colored green and orange in the outer ring, respectively. The number of SRGs found in the Hadza, UHGG, or both is shown as a horizontal line. Hadza genomes that are novel at the family or order level according to GTDB are annotated with red and blue stars, respectively. **(B)** The percentage of bacteriophage species clusters assembled from the Hadza that are novel at the species level according to the MGV ((Nayfach et al., 2021b)), categorized by phylum of the predicted host. Bacteriophages without a host prediction are labeled “Uncharacterized”. **(C)** A phylogenetic tree of eukaryotic genomes recovered from Hadza and Nepali gut metagenomes based on universal single copy genes. Public reference genomes are marked with blue text labels. The heatmap shows the prevalence of the individual eukaryotes in the Hadza, Nepali and Californian cohorts. **(D)** For each population, the percentage of predicted proteins from recovered genomes that are present in the UniRef100 and UHGP-95 (Almeida et al., 2020) protein databases. **(E)** The percentage of metagenomic reads mapping to various domains averaged across all metagenomic samples from each population. The phyla “Bacteriodota” and “Firmicutes_A” are shown separated from other bacteria. “Unmapped” depicts the percentage of reads that do not map to any genomes, and “Low confidence” depicts the percentage of reads that map to genomes with less than 50% genome breadth. **(F)** The Shannon diversity of bacteria, archaea, bacteriophage, and eukaryote genomes in metagenomes sequenced in this study. P-values from two-sided Mann-Whitney-Wilcoxon test with multiple hypothesis correction; *: p < 0.05, **: p < 0.01, ***: p < 0.001, ****: p < 0.0001, ns: p ≥ 0.05. **(G)** Collectors’ curves depicting the average number of genomes detected per sample in each population sequenced in this study after rarefaction to various sequencing depths. The vertical dotted lines indicate the average per-sample sequencing depth of this study (~23 Gbp) and the average depth of samples studied previously (~4 Gbp; ref. (Almeida et al., 2020)). Shaded areas around lines indicate 95% confidence intervals. “Nepal For.” includes the Chepang foragers, while “Nepal Ag.” includes Raute, Raji, and Tharu agrarians.

In parallel to the metagenomic genome recovery efforts described above, which allow study of all microbes in a culture-free manner, we performed anaerobic cultivation and isolation on the same Hadza stool samples. A total of 117 bacterial strains were isolated and subjected to whole-genome sequencing (**Supplementary Table 3)**. These genomes belong to 56 different bacterial species, 18 of which have no previously-cultivated representative and 9 of which are novel relative to UHGG v1 (**Fig. S2**). Phylogenetic analysis shows that genomes we recovered through isolation are highly related to those recovered through metagenomic assembly, corroborating the accuracy of our metagenomic genome recovery pipeline and highlighting the utility of these strains for future laboratory studies of Hadza-associated bacteria.

### Deep Sequencing Reveals Greater Microbiome Insight

The striking level of genomic novelty in the Hadza microbiome uncovered here could be due to the unique lifestyle of the Hadza and/or the exceptionally deep metagenomic sequencing performed **(Fig. 1C)**. In order to compare the Hadza microbiome to populations living other lifestyles without the need for sequence rarefaction, we performed additional deep metagenomic on fecal samples from Nepali and Californian individuals (Jha et al., 2018; Wastyk et al., 2021) **Table S1**). The Nepali samples are from four populations living on a lifestyle gradient: foragers (Chepang), and agrarians (Raute and Raji, recent agrarians; Tharu, longtime agrarians (Jha et al., 2018)). The Nepali and Californian samples were sequenced to the same depth as the Hadza samples, and data were processed using an identical computational pipeline **(Fig. S3)**.

To assess microbiome composition of all samples in light of the newly-recovered genomes from this study, we mapped our metagenomic reads to a series of three custom databases containing full genome sequences of species-level representatives for the bacteria/archaea (n=5,755) bacteriophage (n=16,899), and eukaryote (n=12) genomes (see methods for details). Over 80% of the metagenomic reads from Hadza, Nepali, and Californian samples map to these databases **(Fig. 2E)**. Notably, the Hadza have higher bacterial, bacteriophage, and archaeal diversity than other populations in this study,with the exception of Nepali forager bacteriophage diversity (**Fig. 2F)**. This increased diversity was not due to increased sequencing depth, as an *in-silico* rarefaction analysis revealed more total and novel species of bacteria, archaea, and bacteriophage in Hadza samples compared to other populations across a range of sequencing depths (**Fig. 1G**).

To better understand the impact of sequencing depth across lifestyles we performed an additional *in-silico* rarefaction analysis on the 11 individual samples sequenced to ≥ 50 Gbp, including the deepest publicly-available human gut shotgun metagenome sequenced to date (210 Gbp) **(Fig. 3A)**. The analysis suggests that the Hadza adult gut microbiome contains over 800 bacterial species, compared to ~200 species within Californians sequenced to a similar depths. As deeper sequencing allows detection of lower-abundance microbiome members, we also assessed microbiome composition and genomic novelty of species detected at different abundance levels **(Fig. 3B)**. Rarer species are more likely to be novel across all four populations, even among the Californian cohort. The vast majority of the low-abundance species are Firmicutes, known and common human gut colonists, supporting their status in each population studied as true members of the gut microbiome and not environmental or food-associated microbes. Taken together these data suggest that typically-used shallow sequencing depths present a biased view of the gut microbiome, and that the prevalent use of these depths in the field has led to a systemic bias for highly-abundant species in current genome databases.

**Fig. 3.**
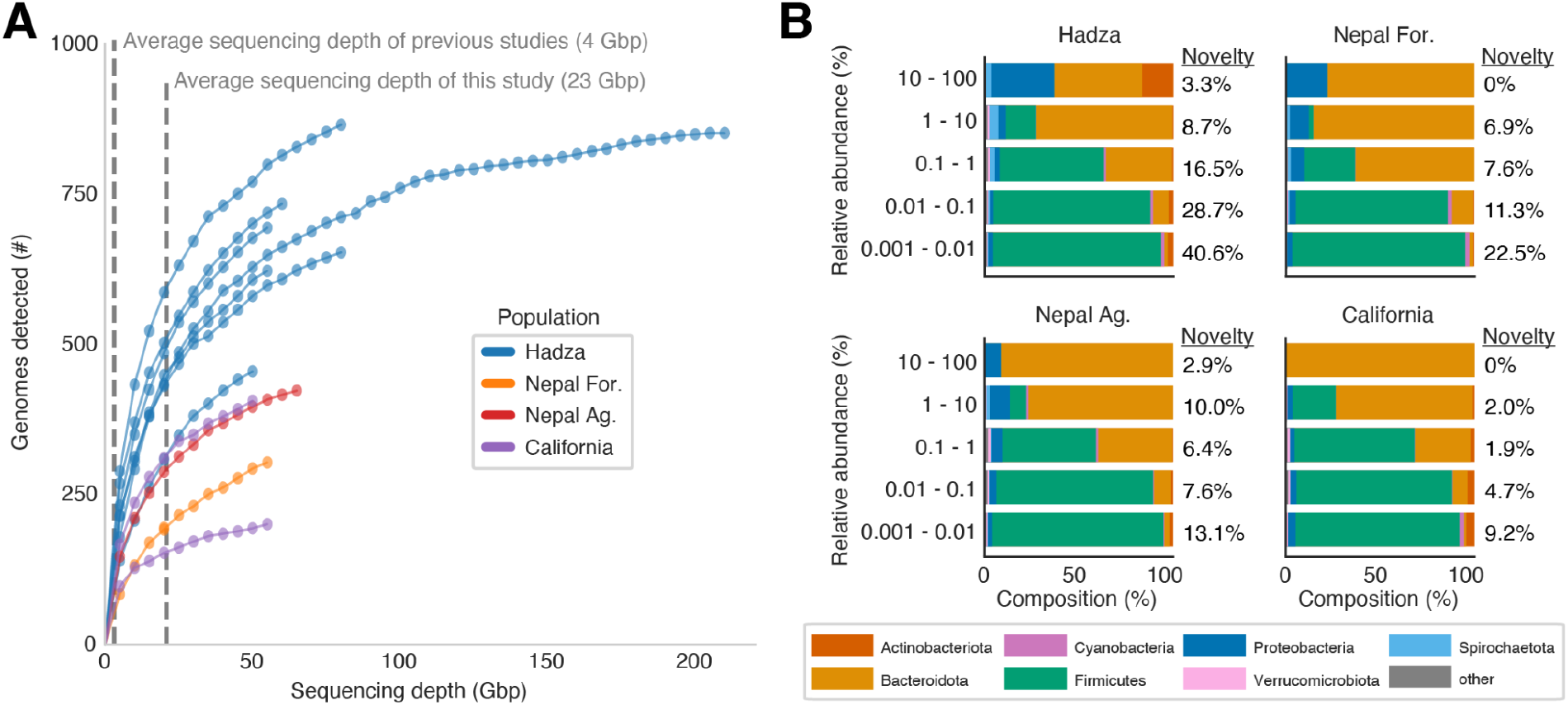
Increased sequencing depth results in the detection of novel and phylogenetically distinct taxa. **(A)** The number of genomes detected in individual samples sequenced in this study when limiting sequencing depth by 5 Gbp increments. Each line represents an individual sample from which ≥ 50 Gbp of trimmed, filtered reads were generated. Lines are colored by population. Vertical dotted lines indicate the average per-sample sequencing depth of this study (23 Gbp) and the average per-sample sequencing depth of samples used in Almleida et al. (4 Gbp (Almeida et al., 2020)). **(B)** Taxonomic distribution of organisms present at different ranges of relative abundance levels (horizontal stacked bar plots) and the percentage of species that are novel according to GTDB r95 (text percentages right of horizontal bars). Organisms detected at low relative abundance levels are more likely to be novel than those that are more abundant.

### VANISH microbes abound in the Hadza

To explore the extent to which the Hadza microbiome differs from other populations, we curated a dataset of 1,800 human gut metagenomes from 21 published studies (Bengtsson-Palme et al., 2015; Brito et al., 2016; Clemente et al., 2015; Conteville et al., 2019; Costea et al., 2017a; Human Microbiome Project Consortium, 2012; Liu et al., 2016; Lloyd-Price et al., 2017; Lokmer et al., 2019; Obregon-Tito et al., 2015; Pasolli et al., 2019; Pehrsson et al., 2016; Qin et al., 2010, 2012; Rampelli et al., 2015; Rosa et al., 2018; Vangay et al., 2018; Zeevi et al., 2015) (industrial, n=950; transitional, n=583; Hadza hunter-gatherers from this study, n=135; and other hunter-gatherers, n=132; **Fig. S4A-B, Table S4)**. Analysis of the hunter-gatherer samples demonstrates that substantial diversity and distinguishing taxa are recovered with deeper sequencing, so subsequent compositional analysis was focused on the deeply sequenced Hadza samples (**Fig. S4C-F**); hunter-gatherer samples from other studies proved difficult to integrate into the analysis due to the shallow depth of sequencing (**Fig. 1A**) and were excluded from the analysis. The presence of each species within our bacterial/archaeal genome database was determined for each sample (**Fig. 4A**, **Table S5**); and VANISH (n=124) and BloSSUM (n=63) taxa were defined as those that are most significantly enriched in the Hadza and industrial populations, respectively (Fisher’s exact test; ≥95th percentile; **Fig. 4B; Fig. S5**).

**Fig. 4.**
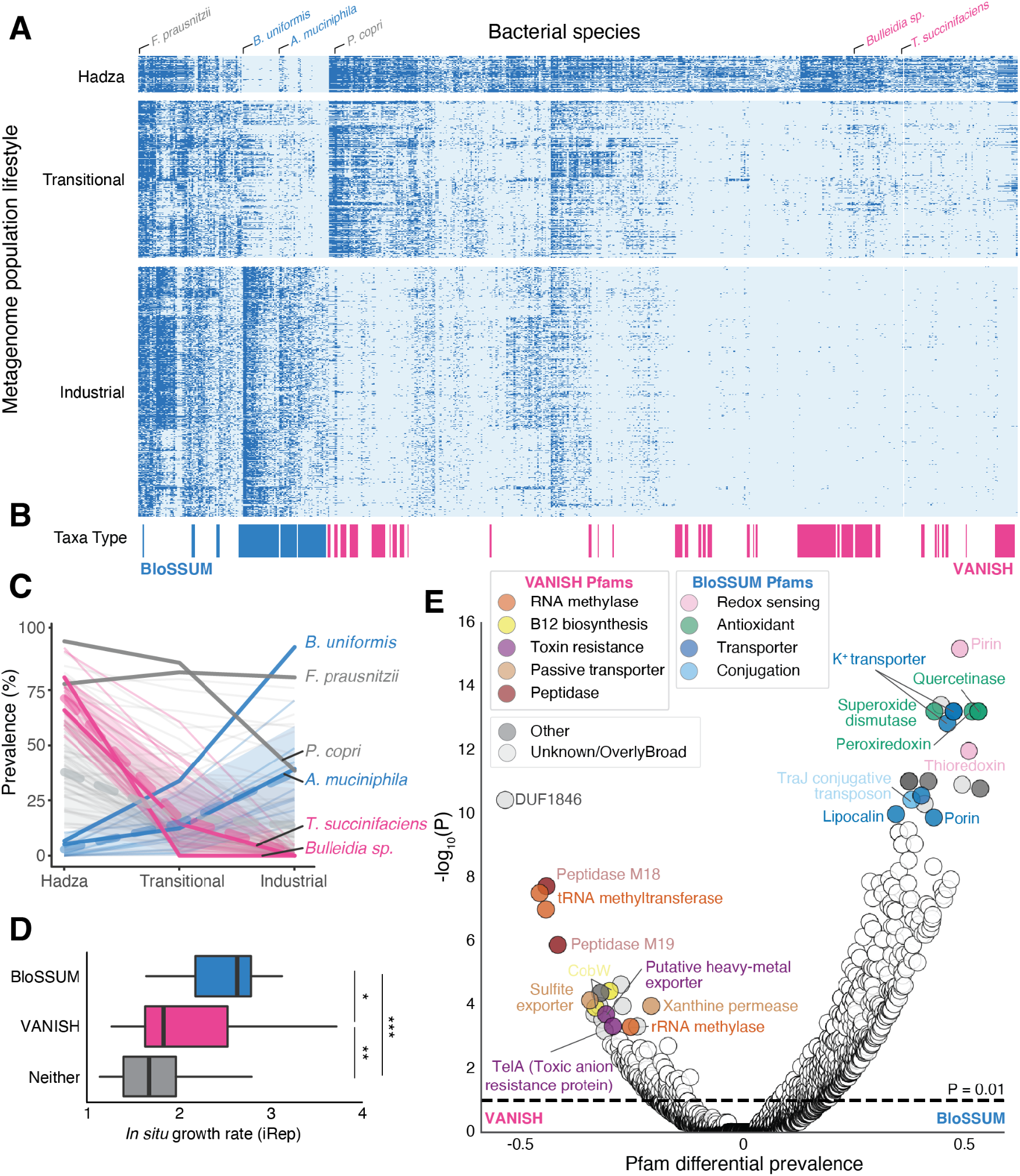
VANISH and BloSSUM taxa have distinct global prevalence, function, growth rates and covariance with eukaryote detection. **(A)** A heatmap depicting the presence of 524 SRGs (columns) within metagenomic samples from populations living different lifestyles (rows). Darker blue indicates SRG presence, lighter blue indicates SRG absence. SRGs with >30% prevalence among all samples in any lifestyle category were included. **(B)** SRGs were classified as “BloSSUM” or “VANISH” based on their prevalence across lifestyles (see methods for details). Colored bars correspond to columns in the heatmap. **(C)** The prevalence of VANISH (magenta), BloSSUM (blue) and non-enriched taxa (gray) in the Hadza, transitional lifestyle populations and industrial lifestyle populations. Dashed lines connect median prevalence across the taxa in each category surrounded by standard deviation (color shaded regions). Solid lines show the median prevalence for 6 representative taxa in each of these lifestyle groups. **(D)** The *in situ* growth rate of SRGs in metagenomes from Nepali individuals, stratified by status as “VANISH” (middle), “BloSSUM” (bottom), or neither (top) (* P ≤ 0.05; ** P ≤ 0.01; Wilcoxon rank-sum test). **(E)** The association of Pfams with VANISH or BloSSUM genomes. The x-axis displays the fraction of BloSSUM genomes a Pfam is detected in minus the fraction of VANISH genomes a Pfam is detected in (Pfam differential prevalence). The y-axis displays the p-value resulting from Fisher’s exact test with multiple hypothesis correction.

Most VANISH taxa (n=120; 96%) and all BloSSUM taxa (n=63; 100%) are detected in “transitional” samples (taken from human populations that have neither hunter-gatherer/forager nor industrialized lifestyles). We find these taxa are typically found at intermediate prevalence, consistent with the extent of lifestyle change corresponding to the magnitude of microbiome shifts (Tamburini et al., 2022) (**Fig. 4C**). Interestingly, BloSSUM taxa have higher *in situ* growth rates than VANISH taxa in transitional samples **(Fig. 4D)** and are negatively associated with the presence of *Blastocystis*, even when comparing within industrialized populations (**Fig. S6**). Replication rate differences may indicate a competitive advantage of BloSSUM taxa over VANISH taxa in the human gut environment.

The observed trade-off between VANISH and BloSSUM taxa concomitant with lifestyle differences poses the question of whether an accompanying trade-off exists with regard to functional capacity in the human gut microbiome. The extraordinary level of novelty present in the Hadza gut precludes the use of most gene annotation pipelines, and we thus focused our analysis on protein domains (Pfams), which represent broad, evolutionary conserved functional units (Mistry et al., 2021). This functional analysis identified 145 and 588 Pfams that are more prevalent in VANISH and BloSSUM taxa, respectively (p < 0.01; Fisher’s exact test, Benjamini p-value correction; **Fig. 4E**; **Table S6**). Pfams most associated with VANISH taxa point to a relatively outsized use of metal ions, peptidases, and RNA methylation. BloSSUM Pfams are associated with antioxidant and redox sensing functionality, perhaps reflecting increased oxygen tension associated with inflammation or an altered epithelial metabolic state in the industrialized gut (Litvak et al., 2018; Sonnenburg and Sonnenburg, 2019b). The difference in the associated functions of enriched Pfams demonstrates that BloSSUM taxa are not functionally redundant to VANISH taxa.

### *Treponema succinifaciens* dispersal mirrors human migration

Several species of the phylum Spirochaetota were identified as VANISH taxa in this study (**Table S5**). Spirochaetota in general, and especially the most well-studied human gut species *Treponema succinifaciens*, are known to be depleted in industrialized microbiomes (Sonnenburg and Sonnenburg, 2019a). Here we leveraged the deep sequencing we performed on Hadza, Nepali, and Californian samples using consistent methods and the 1,047 new Spirochaetota MAGs recovered in this study to conduct a robust analysis of Spirochaetota abundance and prevalence across lifestyles. Our recovered Spirochaetota MAGs belong to the *Treponemataceae*, *Sphaerochaetaceae*, or *Brachyspiraceae* families and span 26 species (including a sequenced isolate of *Treponema perunse (Belkhou et al., 2021)*), 16 of which are novel relative to the UHGG v1. The relative abundance of Spirochaetota species decreases with increased industrialization and no Spirochaetota genomes are detected within Californians **(Fig. 5A)**. Hadza Spirochaetota genomes fall into three diverse families also found in other populations **(colored boxes, Fig. 5B)** suggesting that Spirochaetota are a core component of the non-industrialized lifestyle microbiome and highly susceptible to loss upon lifestyle change.

**Fig. 5.**
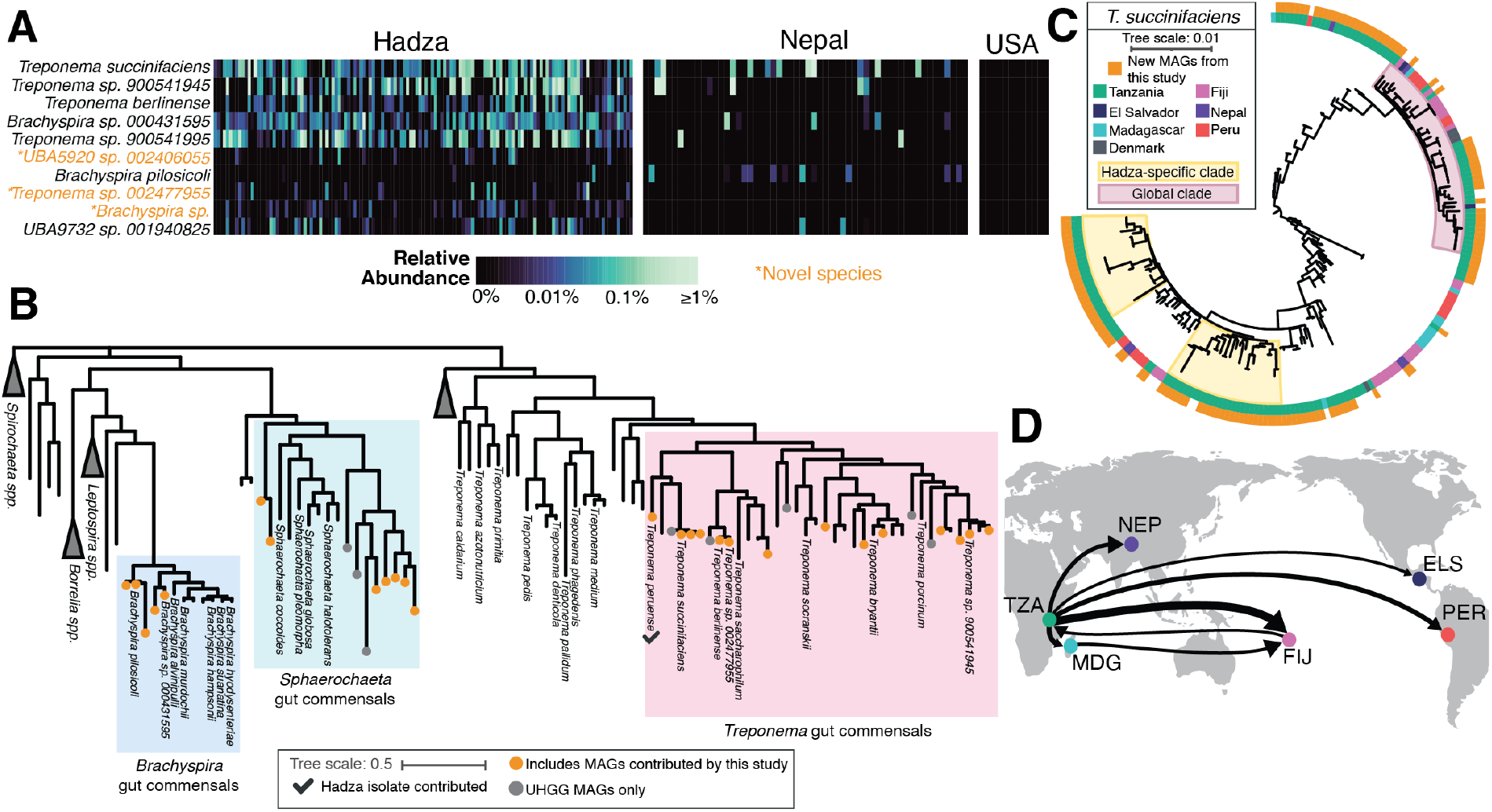
Spirochaetota that are highly abundant in the Hadza are absent in industrial samples. **(A)** A heatmap showing the relative abundance of the 10 most prevalent Spirochaetota species in the Hadza, Nepali, and American cohorts. All samples are sequenced to approximately the same sequencing depth. **(B)** A phylogenetic tree of all Spirochaetota species using genomes from NCBI, the UHGG and the species-representative genomes added in this study. Clades of commensal organisms in the genera *Brachyspira*, *Spirochaeta*, and *Treponema* are highlighted. **(C)** A phylogenetic tree of all *Treponema succinifaciens* MAGs in the UHGG in addition to new MAGs recovered in this study (annotated in outer ring). The inner ring is colored based on the country of origin of the individual contributing the MAG. **(D)** World map showing locations of populations from which *T. succinifaciens* MAGS were recovered as nodes (TZA = Tanzania, MDG = Madagascar, NEP = Nepal, FIJ = Fiji, PER = Peru, ELS = El Salvador). Arrows indicate the detection of transition events between populations as detected by stochastic character mapping. Thickness of the arrow indicates frequency of the transition event (thickest arrow is Tanzania to Fiji, 17.1%). The top 7 most frequent transition events are shown, accounting for 65.7% of all transitions.

The MAGs recovered here increase the number of publicly available *Treponema succinifaciens* genomes from 125 to 346 (276% increase), enabling a robust phylogenomic analysis of the species (**Fig. 5C**). We identified both Hadza-specific and globally distributed clades of *T. succinifaciens* and observed a significant association between phylogeny and continent of origin (p-value<0.0001; delta statistic d=7.79) (Borges et al., 2019). To model the dispersal of *T*. *succinifaciens* between human populations, we performed stochastic character mapping on the phylogenetic tree of MAGs in which the country of origin of each MAG was coded as a trait of the genome and the frequency of “transition events” between each pair of populations is quantified (Suzuki et al., 2021) **(Fig. 5D)**. The 4 most frequent transition events between populations are from the Hadza to other populations, accounting for 46.7% of all transition events, suggesting that *T. succinifaciens* was carried along the out-of-Africa human dispersal routes (Liu et al., 2006). The congruence of *T. succinifaciens* phylogenomics with known patterns of past human migration is consistent with its dispersal being linked to close human contact (e.g., vertical, mother-to-infant, or intergenerational, transmission), as has been described for *Helicobacter pylori* (Falush et al., 2003; Linz et al., 2007).

### Evolution, growth, and dispersal in the Hadza gut

The high sequencing depth and sample number achieved in this study provide an unprecedented opportunity to investigate *in situ* growth rates, microdiversity, and strain sharing within a hunter-gatherer population. To elucidate genes with distinct selective pressures within the Hadza microbiome, we performed an analysis of intra-genic *pN/pS* ratios, a measure of bias towards non-synonymous mutations that suggests positive or diversifying selection (**Fig. 6A**; **Table S6**). Pfams with consistently lower *pN/pS* ratios (p < 0.01; n=520) were often associated with house-keeping annotations, as expected. Notably, however, Pfams with consistently higher *pN/pS* ratios (p < 0.01; n=693) were often associated with extracellular or membrane-bound proteins, such as Ig-like folds, pilin motifs, and collagen-binding proteins. These data provide a roster of functions within the Hadza gut that show relatively increased signatures of diversifying selection, likely in response to dynamics relevant to the Hadza gut such as seasonality of diet, antigen escape from host immune response (Lizano et al., 2007), and phage predation (Rodriguez-Valera et al., 2009).

**Fig. 6.**
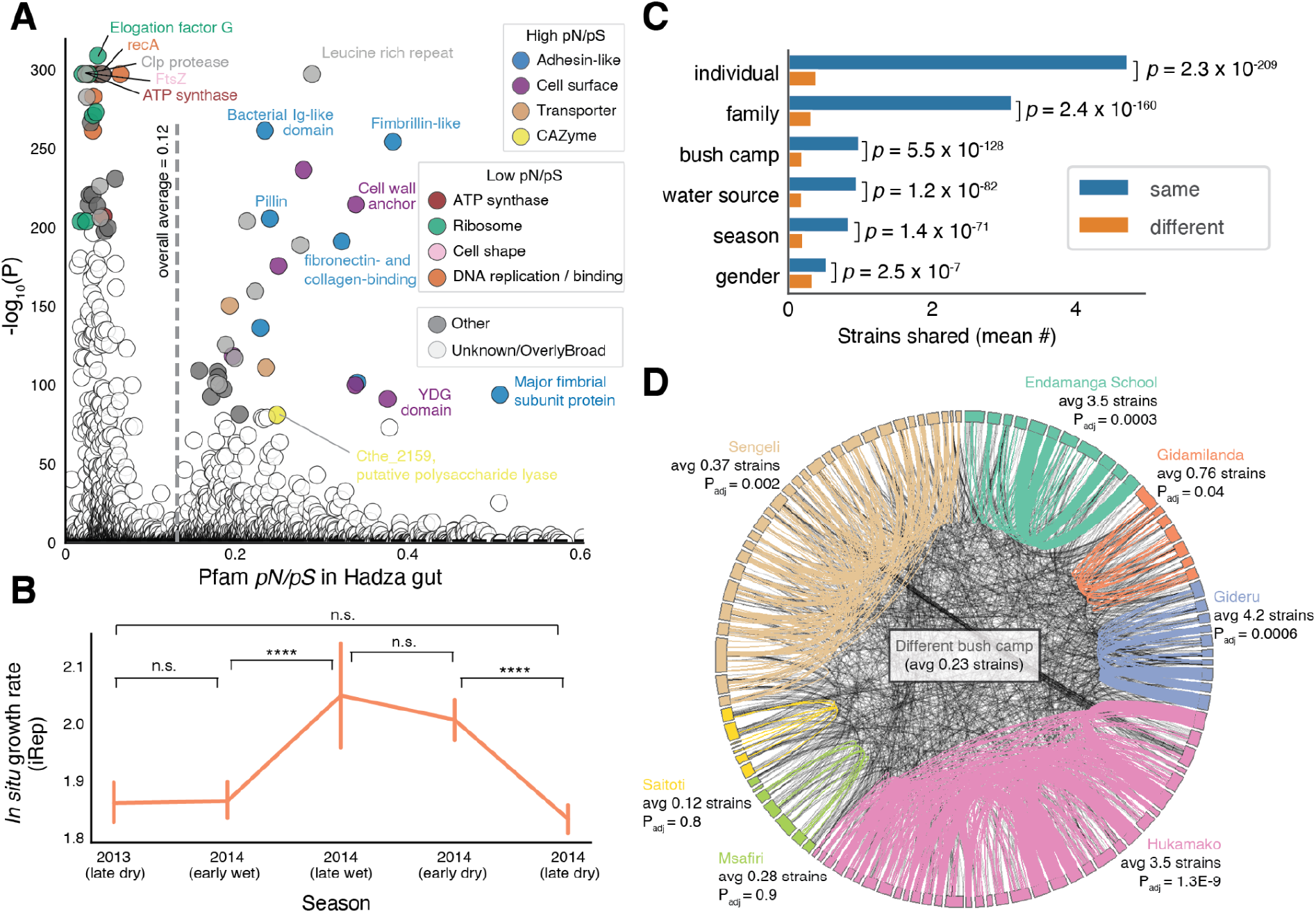
Microdiversity, growth rates, and patterns of strain sharing among Hadza gut bacteria. **(A)** Pfams with high or low *pN/pS* values in Hadza fecal metagenomes. The x-axis displays the mean *pN/pS* value of all genes annotated with each Pfam within Hadza fecal metagenomes. The y-axis displays the probability that the number of times genes annotated as each Pfam were in the top 10% or bottom 10% of all genes on detected genomes was due to random chance (binomial test with multiple hypothesis correction). The 30 Pfams with the lowest p-values for low and high *pN/pS* were manually annotated with broad functional categories. **(B)** *In situ* growth rate measurements of all taxa detected in Hadza adult metagenomes across seasons. Error bars indicate 95% confidence intervals. (n.s. P > 0.05; **** P ≤ 0.0001; Wilcoxon rank-sum test). **(C)** Rectangles along the circumference represent Hadza individuals and each link drawn between boxes indicates a shared strain. Links between members of the same bush camp are colored based on the bush camp; links between bush camps are colored black. The mean number of strains shared between members of the same bush camp and the p-value comparing strains sharing among members of that bush camp vs members from different bush camps are shown (Wilcoxon rank-sums test). **(D)** The mean number of strains shared between Hadza adults broken down by various types of familial relationships. Exact p-values shown from Wilcoxon rank-sum test.

The Hadza gut microbiome has been previously shown to undergo seasonal cycling in carbohydrate-active enzyme (CAZyme) and species composition (Fragiadakis et al., 2019; Smits et al., 2017), and here we confirm these findings using deeper sequencing and updated metagenomic methods **(Fig. S7)**. The deep sequencing performed in this study also allowed us to, for the first time, measure whether *in situ* growth rates exhibit seasonal cycling in the Hadza gut as well. Average bacterial replication rates are lowest in the late dry and early wet seasons, highest in late wet and early dry seasons, and equivalent between the 2013 and 2014 late dry season **(Fig. 6B)**. This pattern of seasonal cycling in the replication rates of Hadza gut bacteria may be driven by i) persistent microbial colonists changing their growth in response to seasonally available foods, or ii) the displacement of bacteria based on seasonally-matched distinct growth strategies.

Family relation and cohabitation are among the strongest factors associated with microbial strain sharing in industrial populations (Faith et al., 2013; Valles-Colomer et al., 2022), but it is unknown whether these patterns hold for hunter-gatherer populations like the Hadza. We performed a high-resolution strain-tracking analysis (threshold for same strain = 99.999% popANI) and found that family members share more recently-transmitted strains than unrelated individuals among the Hadza **(Fig. 6C, Table S7)**. Interestingly, strain sharing among members of the same bush camp approaches that between members of the same family **(Fig. 6C)**, and this effect is stronger in some bush camps **(Fig. 6D**). For example, individuals from the Hukamako camp (pink, bottom right of plot) share more strains with one another than family members share on average across all camps. Drinking water source (e.g., spring, stream, riverbed, etc.) and season (late dry, early dry, early wet, or late wet) have been previously linked to gut microbiome similarity (Jha et al., 2018; Smits et al., 2017), and here we demonstrate that these factors are also linked with the sharing of identical microbial strains **(Fig. 6C)**. Overall, these results point to the importance of environmental factors, kinship, and bush camp membership (a social structure with no equivalent in the industrialized populations) in driving strain dispersal among hunter-gatherers.

## Discussion

The data generated in this study represents a one-of-a-kind collection of human gut microbiome data from one of the last remaining hunter-gatherer populations. The Hadza are a modern people facing challenges related to land dispossession, hunger, and lack of access to education, healthcare and political decision-making, though technologies, food, and medicines from urban centers are becoming increasingly available (Mangola et al., 2022). The data generated from Hadza fecal samples in this study (collected in 2013-14) may thus represent a critical permanent reference point for microbiome scientists to understand the impacts of industrialization on the gut microbiome.

All data generated in this study, and results from all analyses performed in this study, represent a considerable resource to the scientific community considering the large existing gap in general microbiome characterization, including deep metagenomic sequencing in populations living non-industrialized lifestyles. These data and analyses are made freely available to the scientific community. These resources include anonymized metadata, raw metagenomic sequencing reads, full metagenomic assemblies, all MAGs and isolate genome sequences, bacteriophage host identifications, growth rate and population genomic information, millions of genes with UniRef100 annotations, and species-level abundance information across 1,800 public metagenomes across a range of lifestyles. Crucially, the isolation and sequencing of bacterial strains from the same fecal samples on which we performed metagenomics highlight the accuracy and reliability of our computational approach. The genomes recovered in this study will lead to better profiling in future studies, and should encourage the field to adopt deeper sequencing techniques.

In this study we elucidate many novel facets of lifestyle differentiation in the gut microbiome, particularly among the Hadza hunter-gatherers of Tanzania. The discovery of numerous novel clades of bacteria, archaea, bacteriophage, and eukaryotes highlight a leap in understanding of non-industrialized human microbiomes and reframe the incompleteness and bias of commonly used microbial genome reference databases. Functional differences in the gut microbiomes of humans living different lifestyles highlight the consequences of our intestinal inhabitants adapting to a changing gut environment. The VANISH taxa found in present-day Hadza may represent lineages of microbes that shaped human development throughout our species’ long history as foragers. Global phylogenomic analysis of the commensal spirochaete, *Treponema succinifaciens*, shows strain relatedness consistent with known human migration patterns prior to industrialization. Extending deep metagenomic sequencing to populations living across additional geographies will enable a better understanding of which microbes traveled with, were lost, or gained in human populations as we spread around the planet. An important challenge is to characterize the impact of these microbes on human physiology and determine in which contexts the absence or presence of species and functions are beneficial or detrimental to human health. Overall, our results conclusively show that the differences between industrialized and non-industrialized microbiomes go well beyond simple taxonomic membership and diversity. These findings have substantial implications for how the microbiome may be investigated towards improving the health of both industrialized and non-industrialized populations.

## Acknowledgements

We are indebted to the participants in this study. We acknowledge the numerous people and organizations who provided logistical support and conducted sample collection in the USA, Tanzania, and Nepal, including Dorobo Safaris, the Human Food Project, John Changalucha, Alphaxard Manjurano, Maria Gloria Domiguez-Bello, Michelle St. Onge, Allison Weakley, Samuel Smits, Gabriela Fragiadakis, Hannah Wastyk, Yoshina Gautam, Dinesh Bhandari, Sarmila Tandukar, Katharine Ng, Guru Prasad Gautam, Jeevan B. Sherchand, and members of the Gardner lab at Stanford. The sequencing depth and breadth of this study was made possible by the Chan-Zuckerberg Biohub. JLS is a Chan-Zuckerberg Biohub Investigator. The content is solely the responsibility of the authors and does not necessarily represent the official views of the National Institutes of Health.

## Recognition of work with indigenous communities

Research involving indigenous communities is needed for a variety of reasons including to ensure that scientific discoveries and understanding appropriately represent all populations and do not only benefit those living in industrialized nations (Abdill et al., 2022; Green et al., 2020). Special considerations must be made to ensure that this research is conducted ethically and in a non-exploitative manner. In this study, we performed deep metagenomic sequencing on anonymized fecal samples that were collected from Hadza hunter-gatherers in 2013/2014 and were analyzed in previous publications using different methods (Fragiadakis et al., 2019; Smits et al., 2017). The samples were collected legally and ethically with consent from the Tanzanian government and in consultation with Tanzanian field-guides. A material transfer agreement with the National Institute for Medical Research in Tanzania specifies that stool samples collected are used solely for academic purposes, permission for the study was obtained from the National Institute of Medical Research (MR/53i 100/83, NIMR/HQ/R.8a/Vol.IX/1542) and the Tanzania Commission for Science and Technology, and verbal consent was obtained from the Hadza after the study’s intent and scope was described with the help of a translator. The publications that first described these samples included several scientists and Tanzanian and Nepali field-guides as co-authors for the critical roles they played in sample collection, but as no new samples were collected in this study and we do not have a field research program in Tanzania, only scientists who contributed to the analyses described here were included as co-authors in this publication. In July 2022 a collaborative team prepared educational materials translated in the local written language and sent them to Hadzaland with a qualified academic researcher to explain the results of the study.

## Author contributions

Conceptualization: BDM, MMC, MRO, DD, ARJ, EDS, JLS

Methodology: BDM, MMC, MRO, DD, SJ, JLS

Software: BDM, MMC, MRO, DD, SJ

Investigation: BDM, MMC, MRO, DD

Resources: BY, NN, ARJ

Writing - Original Draft: BDM, MMC, MRO, DD, EDS, JLS

Writing - Review & Editing: BDM, MMC, MRO, DD, ARJ, EDS, JLS

Visualization: BDM, MMC, MRO

Funding acquisition: JLS, EDS, MRO, DD

Supervision: EDS, JLS

Project administration: JLS

## Data and materials availability

The authors declare that the data supporting the findings of this study are available within the paper and its supplementary information files. Metagenomic reads and *de novo* genomes are being submitted to the short read archive (SRA) and GenBank and this manuscript will be updated with additional accessions when the submission is complete.

## Materials and methods

### Sample collection

Samples from Tanzania are from 2013-2014 and described previously (Fragiadakis et al., 2019; Smits et al., 2017). Permission was obtained from the National Institute of Medical Research and the Tanzania Commission for Science and Technology. For longitudinal samples, one sample from each individual was marked “high_prority” **(Table S1)** and used as noted in statistical analyses that are not robust to multiple samples from the same individual. Nepal samples were obtained previously (Jha et al., 2018) approved by the Ethical Review Board of the Nepal Health Research Council (NHRC) and the Stanford University Institutional Review Board (IRB). U.S. samples were obtained previously (Wastyk et al., 2021). All human samples were de-identified and collected after receiving informed consent from participants.

### Library preparation and sequencing

Shotgun metagenome sequencing was performed on extracted DNA (MoBio PowerSoil) as described previously (Fragiadakis et al., 2019; Smits et al., 2017). Deeper shotgun metagenome sequencing was performed on samples extracted using phenol:chloroform:isoamyl alcohol described previously (Smits et al., 2017). 101 Hadza individuals were sampled once and 66 individuals were sampled longitudinally. DNA extraction was performed using mechanical extraction (n=318), phenol chloroform extraction (n=38), or both (n=32).

Libraries were prepared using half-reactions (Nextera Flex), using a minimum of 10 ng of DNA and 6 or 8 PCR cycles to minimize amplification bias using a different 12 base pair unique dual-indexed barcode. Libraries were quantified (Agilent Fragment Analyzer) and size-selected (AMPure XP beads,Beckman), targeting a fragment length of 450bp (insert size 350 bp).

Paired-end sequencing (2×140bp) was performed on a NovaSeq 6000 using S4 flow cells at Chan Zuckerberg Biohub (San Francisco, CA, USA). Samples were randomized across runs and sequenced repeatedly until the target depth was reached. Minimum target depth for each sample was 50 million paired-end reads (~14 Gbp) with a subset of samples sequenced to a minimum target depth of 100 million paired-end reads (~28 Gbp). A total of 8,148 giga base pairs (Gbp) of metagenomic data were generated from 388 Hadza metagenomes (range = 0.7 – 210.3 Gbp, mean = 21.0 Gbp, std dev = 14.5 Gbp), 57 Nepali metagenomes (1,794 Gbp total, range = 14.9 – 84.9 Gbp, mean =31.5 Gbp, std dev = 11 Gbp), and 12 California metagenomes (418 Gbp total, range = 25.2 – 56.8 Gbp, mean = 34.8 Gbp, std dev = 9.2 Gbp) for a total of 10.4 Tbp.

### Metagenome quality control and assembly

Raw sequencing reads were demultiplexed and data originating from the same libraries were concatenated prior to analysis. Raw reads were processed using BBtools suite (Bushnell, 2014). Exact duplicate reads (subs=0) were marked (clumpify), adapters and low-quality bases were trimmed (bbduk;trimq=16 minlen=55), trimmed reads were mapped (BBmap) against the human genome (hg19) with masks over regions conserved broadly in eukaryotes, and duplicate reads were removed. FastQC (Andrews, 2010) was used to ensure read quality. BBMerge was used to merge reads that could be joined unambiguously using the recommended settings (rem k=62 extend2=50 ecct vstrict) (Bushnell et al., 2017).

Metagenomes were assembled individually (metaSPAdes (Nurk et al., 2017); v3.13) using unmerged forward/reverse and merged reads (-k 21,33,55,77) with error-correction enabled. Assembly size and contig metrics were evaluated (QUAST (Gurevich et al., 2013) v5.0) and filtered to contigs >=1500 bp for all subsequent analyses. Gene-calling was performed on all assemblies (Prodigal (Hyatt et al., 2010); v2.6.3) in metagenome mode.

### Strain isolation and genome sequencing

Stool resuspended in PBS was plated on CHG, YCFA (Anaerobe Systems), MRS (Sigma Aldrich), BSM (BBL), Colombia (Anaerobe Systems), BHI (Sigma Alrdich), LKV (Anaerobe Systems), *Treponema* media (DSM Medium 275), and milk-enriched media under anaerobic conditions. Individual colonies were re-streaked and then biotyped on a Bruker MALDI-TOF microflex to determine taxonomy. Colonies were grown in liquid media of the same type as the originating agar plate in anaerobic conditions. For isolating *Treponema*, 0.5% agar was added to the liquid media before making plates. Treponema strains were isolated after removing the top layer of agar to harvest colonies within the agar. Many of these isolated strains are not currently amenable to freezer storage and liquid-cultivation-based propagation in isolation.

Genomic DNA was extracted (Qiagen DNeasy Blood and Tissue). Libraries were prepared using half-reactions of the Nextera Flex kit, a minimum of 10 ng of DNA as input, 6 or 8 PCR cycles to minimize PCR amplification bias and a different 12 base pair unique dual-indexed barcode. Libraries were quantified (Agilent Fragment Analyzer) and size-selected (AMPure XP beads; Beckman), targeting a fragment length of 450bp (insert size of 350 bp). Paired-end sequencing (2×140bp) was performed on a NovaSeq 6000 using S4 flow cells at Chan Zuckerberg Biohub. Assembly of genomes was performed by trimming using BBduk (trimq=30), normalizing read depth using BBnorm (target=320, min=2), and assembled using SPAdes v3.13.1 (-k 21,33,55,77,99,127) (Prjibelski et al., 2020). Genomes were assessed for completeness and contamination using CheckM v1.1.2 (Parks et al., 2015).

### Bacterial and archaeal genome recovery and refinement

A novel “co-mapping” approach was developed to leverage contig depth information from multiple, closely related samples and improve genome bin recovery from single-sample assemblies. MASH sketches (-s 1000000 -k 32 -m 2)(Ondov et al., 2016) were created from reads in each metagenome individually, and sketches were compared in a pairwise manner. For each assembly, reads from that sample and the nine next-closest related samples by MASH distance were mapped (Bowtie2 (Langmead and Salzberg, 2012); --very-sensitive -X 1000) and genome bins generated using contig depth for all 10 samples (MetaBAT2 (Kang et al., 2019); v2.15, default settings). For California samples, only samples taken from the same individual were co-mapped. Genome bin quality was assessed using CheckM v1.1.2 (Parks et al., 2015) and anvi’o (Eren et al., 2015) (v6.3).

Bins were refined using MAGpurify v2 (Nayfach et al., 2019) (using weighted mode for gc_content, tetra_freq, and coverage). The database used by Nayfach et al.(Nayfach et al., 2019) for conspecific analysis was augmented by adding all bins that were >=95% complete and <=5% contaminated (CheckM and anvi’o). For each species-level group, only the highest-quality genome bin for each individual was included. Flagged contigs were removed. Rarely, a module suggested the removal of >25% of a bin’s length, and in such cases that module was turned off. Genomes with ≥50% completeness and <10% contamination according to CheckM were retained, in accordance with MIMAG standards (Bowers et al., 2017).

### Evaluation of self- and co-mapping relative to isolate genomes

Isolate genomes from Hadza stool samples were de-replicated (dRep v3.2.2; -s 100000, -sa 0.99). The highest scoring isolate as the representative when multiple isolates from the same secondary cluster were isolated from the same sample. 19 representative isolates were identified from samples that also had metagenome sequencing, assembly, and binning. Representative isolates and bins (>=50% complete, <5% contamination) generated using self-mapping and co-mapping were compared (MASH; -s 100000), selecting most similar bin MASH distance <0.05), with co-mapping and self-mapping recovered 17 and 10 bins representing isolates, respectively, with no significant differences in quality.

### Creating bacteria / archaeal species-level genome database

Bacterial and archaeal genomes sharing ≥ 95% average nucleotide identity (ANI) over 30% of their length were considered the same species (Olm et al., 2020). Species-level groups were determined using dRep (v3.0.0 (Olm et al., 2017);--S_algorithm fastANI --multiround_primary_clustering --clusterAlg greedy -ms 10000 -pa 0.9 -sa 0.95 -nc 0.30 -cm larger) based on the ANI between all genomes within each species-level group. Each genome was assigned a “centrality” score according to its average ANI to all other genomes in the group. The highest score genome was chosen as representative for each species-level group using the formula: score = (1*completeness) – (5*contamination) + (0.5*log10(ctg_N50)) + (1*log10(contig_bp)) + (2*(centrality-0.95)*100).

Centrality was calculated between all genomes in the UHGG genome database (v1.0) using the species-grouping (Almeida et al., 2020), and species representatives were chosen as above. Representatives from *de novo* genomes generated here and from the UHGG database (v1.0) were compared (dRep; --S_algorithm fastANI --multiround_primary_clustering --clusterAlg greedy -ms 10000 -pa 0.9 -sa 0.95 -nc 0.30 -cm larger). Representatives for each species-level group were chosen using the formula: score = (1*completeness) – (5*contamination) + (0.5*log10(ctg_N50)) + (1*log10(contig_bp)). Representatives were compared using the same dRep command, and winners were chosen using the same scoring criteria. Species-level group membership was back-propagated to the original bins.

### Annotating bacteria / archaeal genomes and assessing genomic novelty

Taxonomy was determined for all species-level representative genomes using GTDB (r95) (Chaumeil et al., 2019). Novelty against UHGG v1 was determined based on the species-level clustering described above. Only genomes that pass both the MIMAG genomic standards used in this study (≥50% completeness and <10% contamination) and the standard used during UHGG creation (completeness − (5*contamination) > 50) were considered in comparisons against UHGG. Species groups containing only genomes recovered from the Hadza were considered novel relative to UHGG.

A phylogenetic tree was made (GtoTree (v1.5.36) (Lee, 2019) with bacterial gene sets (-H Bacteria). All other settings were default. The tree was visualized using iTol (Letunic and Bork, 2007) with taxonomy provided by GTDB.

### Eukaryotic genome recovery and analysis

EukRep (v0.6.6) (West et al., 2018) was employed on all assemblies (default settings) and if a genome bin was both >5 mega base pairs and >80% eukaryotic according to EukRep, it was called eukaryotic. EukCC (v1.1) (Saary et al., 2020) was run on eukaryotic bigs using database eukcc_db_20191023_1

Proteins identified via EukCC were compared against UniRef100 (Suzek et al., 2007) (downloaded 3/5/2020) using DIAMOND (Buchfink et al., 2015) with a maximum e-value of 0.0001 (blastp -f 6 -e 0.0001 -k 1). The resulting taxonomy was parsed with tRep (https://github.com/MrOlm/tRep/tree/master/bin) (Olm et al., 2019b). Eukaryotic genomes with the same species-level taxonomy that originated from the same metagenomic sample were presumed to be from the same organism, were merged into a single file and re-analyzed using EukCC and tRep.

Phylogenetic tree was created (GToTree; v1.5.36) (Lee, 2019) “GToTree -H Universal_Hug_et_al -j 4 -B -c 1 -t”) with a custom set of public reference genomes. Tree was visualized using iTol (Letunic and Bork, 2007).

### Creating eukaryotic species-level genome database

To identify eukaryotic species that may be present in the metagenomics sequenced in this study and which did not have genomes recovered using the pipeline described above, we ran the program EukDetect (Lind and Pollard, 2021) on all metagenomes sequenced in this study. Five species were detected in at least two samples with “perecent_observed_markers” ≥ 50, and reference genomes for these five species were included in the eukaryotic species-level genome database. In addition to these five genomes, the highest quality representative genome from each of the seven species of eukaryotes recovered in this study was included in the eukaryotic species-level genome database.

Metagenome reads were mapped onto the eukaryotic species-level genome database (Bowtie 2 (Langmead and Salzberg, 2012)) and the resulting mappings were processed (inStrain quick_profile; v1.2.14 (Olm et al., 2021) and CoverM v0.4.0 (https://github.com/wwood/CoverM)). A species was “present” if the breadth of coverage according to inStrain exceeded 0.1.

### Viral genome recovery

CheckV (Nayfach et al., 2021a) (version 0.8.1, end-to-end mode, database v1.0) was run on all assembled contigs >=1500bp. Contigs predicted to contain one or more proviruses were run iteratively through CheckV (up to 5 rounds) until CheckV assumed the remaining region was viral. For provirus iterations only yielding an HMM-based completeness estimates, the most complete fragment was selected and excised from the parent contig. For provirus iterations with AAI (Average Amino acid Identity)-based completeness predictions, the fragment with the length closest to expected length was selected and excised from the parent contig. Viral contigs were passed through the MGV viral detection pipeline (Nayfach et al., 2021b) and Bacphlip (v0.9.6) was run to assign a lytic and temperate score (Hockenberry and Wilke, 2021).

### Creating bacteriophage species-level genome database

The 40,171 viruses recovered in this study were clustered into species-level groups as described previously (Nayfach et al., 2021b) (blastn --min_ani 95 --min_qcov 0 --min_tcov 85, https://github.com/snayfach/MGV/tree/master/ani_cluster), and the longest viral contig in each cluster was selected as the representative. To measure novelty versus MGV, the 16,899 species-level representatives were subsequently clustered with the 54,118 MGV cluster representatives into species-level groups using the same method, and clusters without an MGV genome were considered novel.

### Viral host prediction

Host prediction was performed on the 40,171 viruses as described previously (Nayfach et al., 2021b). Briefly, CRISPR spacers were identified (PILER-CR (Edgar, 2007) and CRT (Bland et al., 2007)). BLASTN (Camacho et al., 2009) was used to search viruses for CRISPR spacers identified from bins reported here and UHGG v1 (-dust no -word_size 18). CRISPR spacer hits were retained if there was a maximum of one mismatch or gap over >=95% of the spacer length. Additionally, hs-blastn (Chen et al., 2015) was used to identify >=1kb and >=96% DNA identity hits between all UHGG and newly-recovered genomes and viruses reported here. All viral connections to host genomes were aggregated, and host taxonomy was assigned based on the lowest host taxonomic rank that had >70% agreement across CRISPR or BLASTN.

### Characterizing diversity

Reads from all metagenomes generated here were mapped to the bacterial/archaeal, bacteriophage, and eukaryote species-level genome databases (Bowtie 2 (Langmead and Salzberg, 2012)). Resulting mappings were processed (inStrain quick_profile; v1.2.14 (Olm et al., 2021) and CoverM v0.4.0 (https://github.com/wwood/CoverM). Prokaryotes where the representative genome was detected at ≥ 0.5 breadth (i.e. at least half of bases were covered by at least 1 read) were considered present. Bacteriophages and eukaryotes breadth thresholds were 0.75 and 0.1, respectively.

Relative abundance (% DNA) was calculated as (# reads mapping a genome / total # reads in metagenome). Shannon diversity was calculated based on relative abundance (% DNA) values (scikit-bio (http://scikit-bio.org)).

### Rarefaction analysis

*In silico* rarefaction was performed on samples sequenced to ≥ 50 Gbp using the InStrain auxiliary script “rarefaction_curve.py” (v0.3.0) (https://github.com/MrOlm/inStrain/blob/master/auxiliary_scripts/rarefaction_curve.py) on a .bam file of reads mapped with Bowtie 2 (Langmead and Salzberg, 2012). For other rarefaction curves **(Fig. 2)** an alternative *in silico* rarefaction technique was used. Genomes with < 50% breadth were removed from the analysis, and for each rarefaction level 1) a scaling threshold was established based on the total sequencing depth (scaling factor = rarefaction depth / total sequencing depth), 2) scaled genome coverage was calculated by each genome by multiplying un-rarefied coverage by this scaling factor, and 3) genomes with scaled coverage ≥ 1 were considered detected.

### Collating previously published human gut metagenomic samples

Prevalence of microbial species across lifestyle was characterized using a curated collection of 2122 metagenomes including samples from industrial (Bäckhed et al., 2015; Bengtsson-Palme et al., 2015; Human Microbiome Project Consortium, 2012; Lloyd-Price et al., 2017; Pehrsson et al., 2016; Qin et al., 2010, 2012; Zeevi et al., 2015), transitional (Brito et al., 2016; Costea et al., 2017b; Liu et al., 2017; Lokmer et al., 2019; Pehrsson et al., 2016; Rosa et al., 2018; Tett et al., 2019; Vangay et al., 2018), and hunter-gatherer populations (Conteville et al., 2019; Lokmer et al., 2019; Pasolli et al., 2019; Rampelli et al., 2015). Samples were binned using the U.N. Human Development Index (HDI) (Groussin et al., 2021). Samples from individuals < 3 years old were excluded. For longitudinal samples, a single sample was randomly selected resulting in 137 Hadza samples. Reads were processed as described above. Samples with fewer than 60 genomes detected were excluded.

Hadza sample ERR7803603, sequenced to a depth of 210 Gbp, was determined to be the deepest human gut metagenome sequenced as of 28 Feb 2022 by downloading all summary metadata from NCBI SRA with the search term “(txid408170[Organism:noexp]) AND WGS[Strategy]” and sorting by decreasing base pairs sequenced.

### Species prevalence analysis

All reads generated here and publicly available were mapped to the bacterial/archaeal species-level genome database (Bowtie2 (Langmead and Salzberg, 2012)), and resulting mappings were processed using inStrain quick_profile (v1.2.14) (Olm et al., 2021) and CoverM v0.4.0 (https://github.com/wwood/CoverM)). Species detected at ≥ 0.5 breadth were considered present and prevalence was calculated as the percentage of metagenomes in which the species was present.

Genomes were assigned to VANISH or BloSSUM using p-values resulting from Fisher’s exact test on the following contingency table: [[(# Hadza samples where genome is found, # industrial samples where genome is found), (# Hadza samples genome is not found, # industrial samples where genome is not found)]]. All p-values were ranked and a percentile score was assigned. Genomes in the 95th percentile or greater where Hadza prevalence was higher were “VANISH” taxa. Those in the 95th percentile or greater where industrial prevalence was higher were “BloSSUM”.

Heatmaps displaying species prevalence data were created using the R package “pheatmap” (v1.0.12). Principal coordinate analysis was performed on the species prevalence data using the vegdist function in the package “vegan” (Dixon, 2003) (v2.5-6) and the function cmdscale from the package “stats” (v4.0.4).

### Growth rate analyses

InStrain profile (v1.2.14) (26) was run on all .bam files created as described in the “species prevalence analysis” section. All iRep values for genomes with ≥ 50% genome breadth and with values < 5 were considered valid. Seasonality of iRep values was plotted using seaborn v0.11.1 (Waskom, 2021) “lineplot” with the default estimator (mean) and 95% confidence interval for error bars.

### Blastocystis analysis

Presence or absence of each *Blastocystis* MAG was determined as described above. The top two most prevalent *Blastocystis* MAGs were most closely related to *Blastocystis ST1* and *Blastocystis ST4*, respectively (tRep; https://github.com/MrOlm/tRep/tree/master/bin) (Olm et al., 2019b). Wilcoxon rank sum test was used to determine if presence of a *Blastocystis* genome was correlated with total relative abundance of VANISH taxa and BloSSUM taxa separately. Linear discriminant analysis was performed using the “lda” function from the package MASS (v7.3) to determine the effect size of each association.

### Seasonality analysis

Principal coordinate analysis was performed on the Bray-Curtis distance between all Hadza samples in our study. Relative abundance was aggregated at the taxonomic level of family to mirror initial analysis done in Smits, et al. (Smits et al., 2017). The adonis function in the R package “vegan” was used to test significance by season. Subject ID was used as a sub-stratum. A Wilcoxon rank-sum test was used to determine whether samples varied in composition along the major axis of variation, aggregated by season.

The average relative abundance of each species-level group in our bacterial/archaeal species-level genome database was calculated for each sub-season. Taxa that observed cyclical abundance over the course of a year was determined (Kruskal-Wallis test; p-values were Bonferroni-adjusted to control for multiple hypothesis testing).

CAZyme annotation was performed using dbCAN_v9 HMMs (Zhang et al., 2018) (http://bcb.unl.edu/dbCAN2/download/Databases/V9/dbCAN-HMMdb-V9.txt). Proteins were searched against the HMM collection using hmmscan (Eddy, 2011) and filtered using the “hmmscan-parser.sh” script provided with dbCAN2. Seasonal CAZyme analysis was performed using previously described seasonal delineations (Smits et al., 2017).

### Protein clustering and novelty assessment

Predicted proteins were clustered at 95% identity (MMseqs2 (Steinegger and Söding, 2017); v12.113e3; easy-linclust --cov-mode 1 -c 0.8 --kmer-per-seq 80 --min-seq-id 0.95 --compressed 1). Novelty relative to UHGP-95 (v1.0) (Almeida et al., 2020) was determined by clustering together UHGP-95 with our *de novo* representative proteins (MMseqs2) and back-propagating to the initial *de novo* clustering to calculate the number of protein clusters assembled from each lifestyle. Representative proteins were also compared against UniRef100 using DIAMOND (Buchfink et al., 2015). Novel proteins were defined when the representative protein was not related to any protein in the UniRef100 database with ≥ 95% amino acid identity.

### Protein annotation

Proteins were annotated (Pfams (v32) (El-Gebali et al., 2018); hmmsearch (Eddy, 2011)), filtered (hmmsearch --cut_ga --domtblout), and protein domain overlap was resolved (cath-resolve-hits.ubuntu14.04 (Lewis et al., 2019); --input-format hmmer_domtblout --hits-text-to-file).

### Pfam enrichment analysis

For each Pfam, the number of VANISH and BloSSUM genomes with at least one gene containing a Pfam was recorded as “c1” and “c2”, respectively. Pfams found more often in one genome set or the other were detected using a Fisher’s exact test on the following contingency table: [[c1, (# VANISH genomes) − c1], [c2, (# BloSSUM genomes) − c2]]. Multiple hypothesis correction was performed using the FDR method(Seabold and Perktold, 2010)). Pfam differential prevalence was calculated as (c2 / (# BloSSUM genomes)) − (c1 / (# VANISH genomes)).

### Spirochaetota analysis

Spirochaetota genomes from the bacterial/archaeal species-level genome database and NCBI were de-replicated at the species level (dRep; --S_algorithm fastANI -ms 10000 -pa 0.9 -sa 0.95), and a phylogenetic tree was generated (GtoTree;v1.5.36) (Eddy, 2011; Edgar, 2021; Gutierrez et al., 2009; Hyatt et al., 2010; Lee, 2019; Tange, 2018)) from bacterial (-H Bacteria) gene sets. All other settings were default. The tree was visualized using iTol (Letunic and Bork, 2007) and colored by taxonomy provided by GTDB.

A phylogenomic tree of *Treponema succinifaciens* in the bacterial/archaeal species-level genome database was generated using GToTree; v1.5.36) (Eddy, 2011; Edgar, 2021; Gutierrez et al., 2009; Hyatt et al., 2010; Lee, 2019; Tange, 2018) with IQ-Tree (Minh et al., 2020) from bacterial (-H Bacteria) gene sets (completeness threshold 75% with “-G 0.75”). We used country-of-origin information (re-coded as continent-of-origin) as a trait of each genome to measure the degree of phylogenetic signal in the geographic spread of the MAGs (“delta” function from Borges, et al. (Borges et al., 2019)). *P*-value of the delta statistic was performed using 100 calculations with randomly permuted tree tip labels.

### Stochastic character mapping of *Treponema succinifaciens*

Stochastic character mapping was performed using SIMMAP via the “make.simmap” function (“phytools” R package (Revell, 2012)). We applied the character mapping on the marker-based tree of *T. succinifaciens* GToTree generated MAGs (described above). “Country of origin” of each MAG served as a trait and inferred ancestral character states on phylogeny (equal rates model, repeated 100 times to calculate average # of character changes and direction of host transfer events).

### Pfam pN/pS analysis

The *pN/pS* was calculated using inStrain (v1.2.14) (inStrain profile --database_mode) (Olm et al., 2021) on mappings to the bacterial/archaeal species-level genome database, using the predicted genes. All genomes detected with < 80% breadth were excluded from analysis. For remaining genomes, genes with “SNV_count” < 5 were excluded. If <10 genes in a genome fit this criteria, the genome was excluded. Genes with ≥ 5 “SNV_count” and a blank “pNpS_variants” value were assigned a “pNpS_variants” of 100. Genes were sorted according to “pNpS_variants”, and genes in the top and bottom 10% of “pNpS_variants” were recorded. How many times each Pfam was detected on any genes that passed the above filters (“trial_count”) and how many times the Pfam was in genes in the top and bottom 10% of genes based on “pNpS_variants” (“top_success_count”, “bottom_success_count”) was noted.

To determine Pfams in the top or bottom 10% of “pNpS_variants” more often than expected by chance, genes detected in less than 5 samples were excluded, the number of times a gene was in the top 10%, bottom 10%, and seen total was scaled (“trial_count”/5), and the scaled “top_success_count”, “bottom_success_count”, and “trail_count” values were summed together. Probability that the “top_success_count” or bottom_success_count” was due to random chance was calculated using binomial statistics (Python Scipy(Jones et al., 2001)). P-values reported as 0 were set to 1E-300 and multiple hypothesis correction was performed (FDR (Seabold and Perktold, 2010)). Mean Pfam pN/pS was calculated as the average “pNpS_variants’’ of all genes on genomes with ≥ 80% breadth and a non-blank “pNpS_variants” value.

The procedure described above was repeated using “coverage” instead of “pNpS_variants” to detect Pfams associated with genes with higher or lower coverage than others. To avoid mis-mapping (recruiting genes from other populations), all Pfams with uncorrected p-values < 0.01 were excluded from the “pNpS_variants” analysis.

### Strain sharing analysis

Genome detection was defined as minCov breadth ≥0.5 (i.e. at least half of bases were covered by at least 5 reads) as measured using “inStrain profile”. Each species detected in more than one individual was compared using inStrain compare. Where a genome was detected in more than 120 samples, samples were divided into groups of equal size such that no group had more than 120 samples, and “inStrain compare” was then run on each group. A distance matrix was created for each species based on resulting popANI values and used to cluster each species into individual strains using “average” hierarchical clustering with a threshold of 99.999% popANI (Scipy cluster). Strains shared between sample pairs were calculated based on this strain definition, and P-values were calculated only considering pairs of samples in which both samples were from Hadza adults.

## Supplemental Information

**Supplementary Table 1**: Description of Hadza, Nepali, and Californian cohorts

**Supplementary Table 2**: Comprehensive genome information info (including representative genomes and other genomes)

**Supplementary Table 3**: Roster of strains isolated from Hadza stool (including cultivation information)

**Supplementary Table 4**: Global metagenomics data set broken down by sample

**Supplementary Table 5**: Prevalence/abundance data for each species-level representative genome in our bacterial/archaeal species-level genome database

**Supplementary Table 6**: Pfam info (lifestyle-enrichment and pN/pS data)

**Supplementary Table 7**: Strain sharing data between Hadza adult samples

---

**Figure S1.**
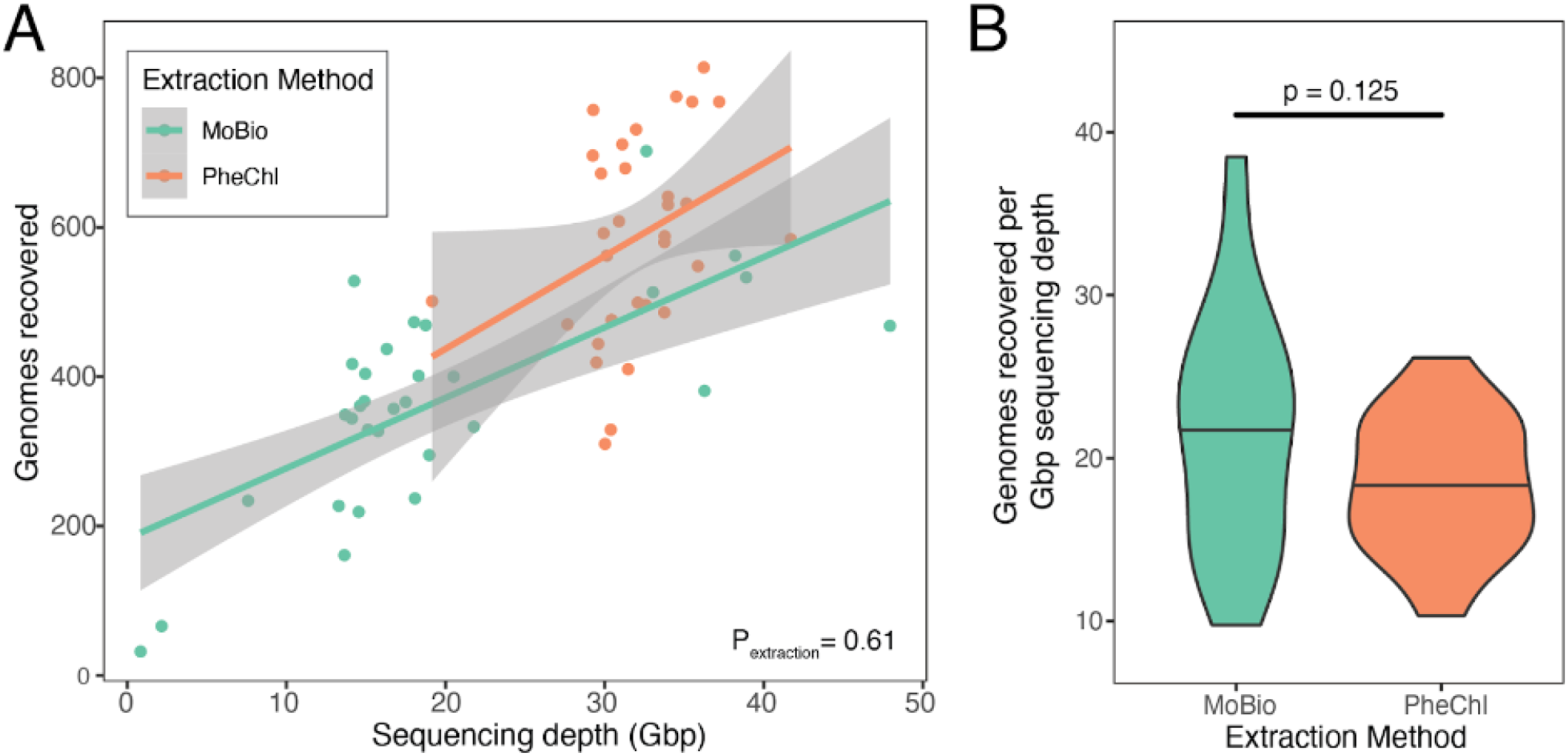
Comparing genome recovery from MoBio extraction vs. phenol chloroform (PheChl) extraction. (**A**) Scatter plot of genomes recovered versus sequencing depth for Hadza samples extracted via both MoBio and PheChl extraction methods. Specifying a linear regression of genomes recovered against sequencing depth with an interaction term for extraction method reveals a non-significant interaction effect (P = 0.61). (**B**) Normalized genomes recovered per Gbp sequencing depth show no statistical difference for samples extracted with both MoBio or PheChl (P = 0.125, Wilcoxon rank-sum test).

**Figure S2.**
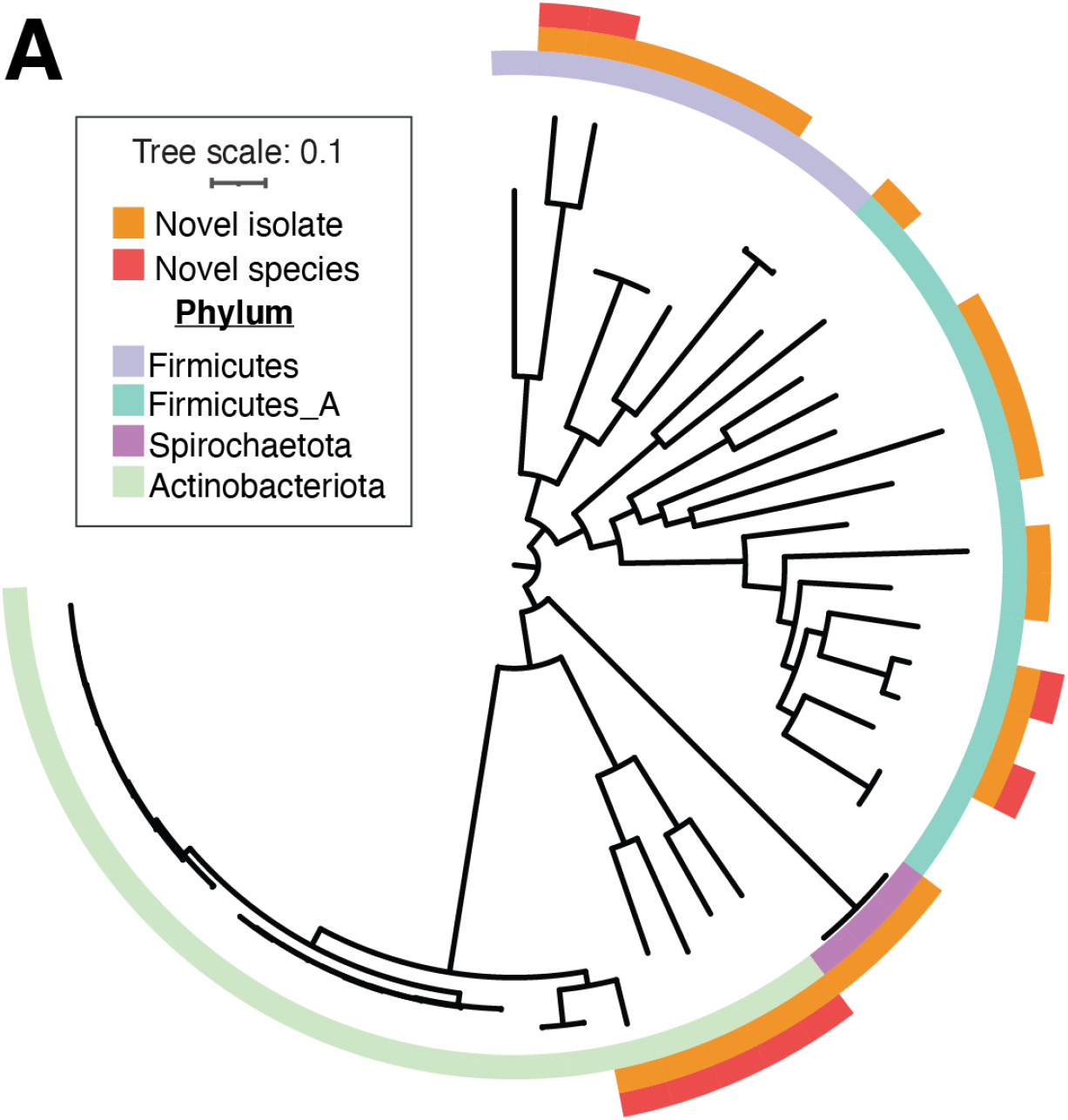
Phylogenies of strains isolated from Hadza stool samples. (**A**) A phylogenetic tree of all isolate genomes sequenced in this study. The tree is decorated with phylum of each species (inner ring), whether the species is newly isolated for the first time (middle ring) and whether the species is novel relative to UHGG (outer ring).

**Figure S3.**
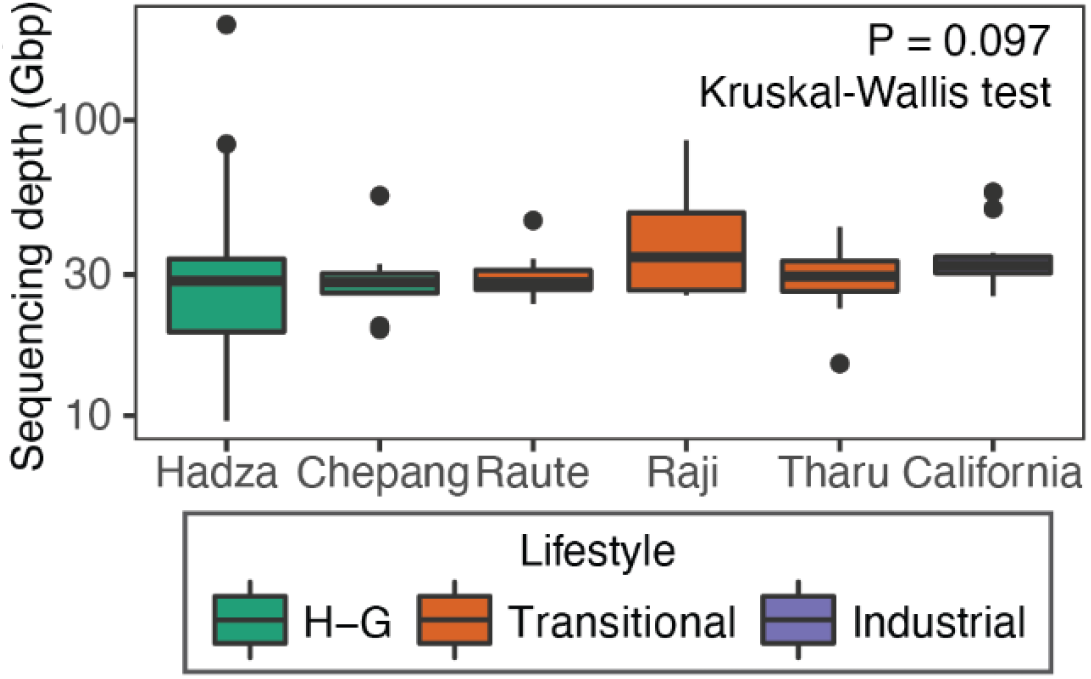
The median metagenomic sequencing depths of populations sequenced in this study. A box plot showing the distribution of sequencing depth, in giga base pairs (Gbp) for each of the populations sequenced in this study. The Chepang foragers and Raute, Raji, and Tharu agrarians are the Nepali populations. The populations do not differ significantly by sequencing depth (P = 0.097, Kruskal-Wallis test).

**Figure S4.**
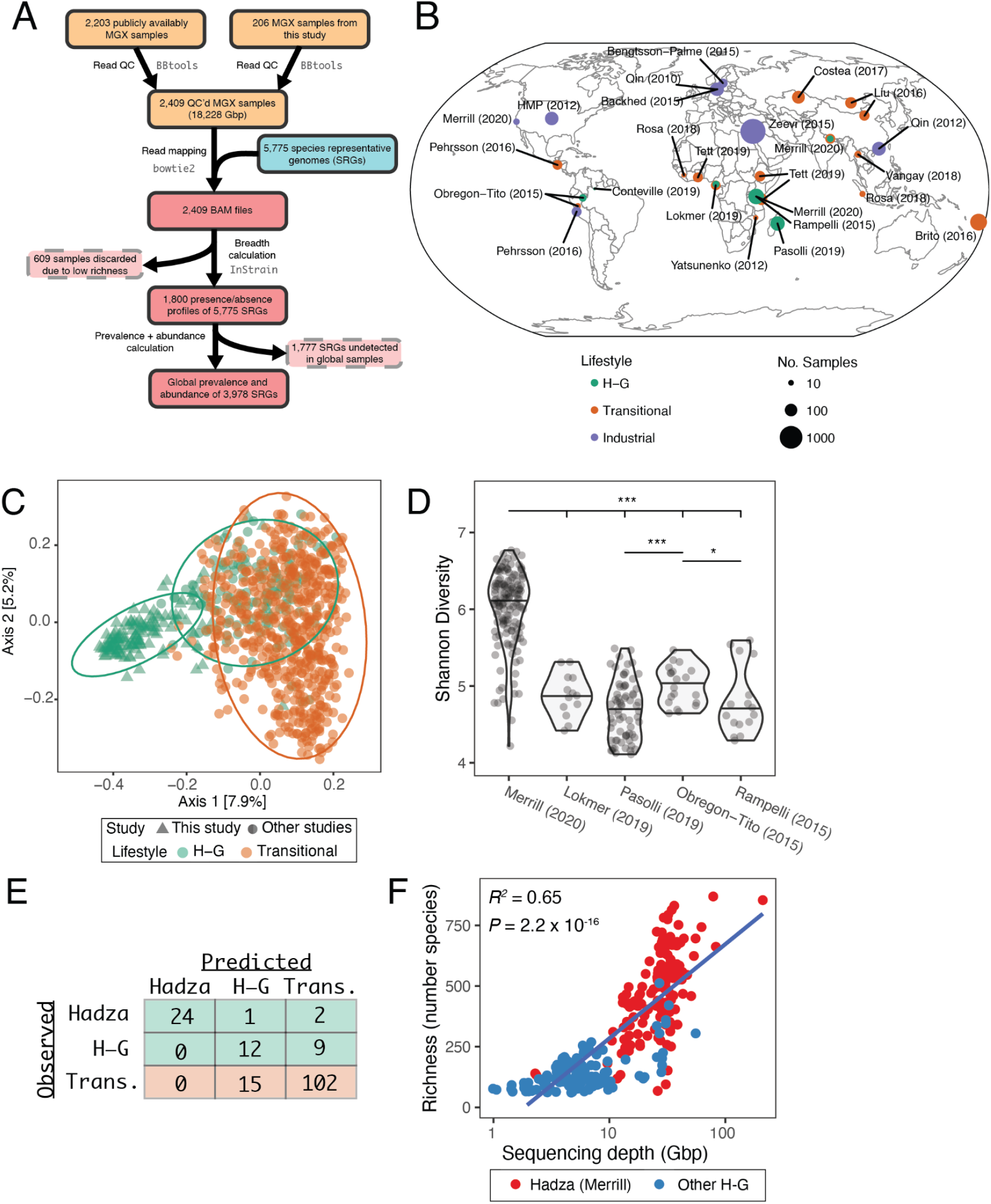
Global metagenomics data set and analysis of publicly available hunter-gatherer samples. **(A)** A flowchart showing the computational pipeline used to analyze global metagenomics samples. **(B)** A world map showing the geographic locations of global metagenomics samples. Dots are colored based on the lifestyle of the study population and the size of the dots indicate the number of samples contributed by that population. ‘H-G’, Hunter-Gatherer. (**C**) PCoA plot of H-G (green) and transitional (orange) samples in our global metagenomics data set. Triangles are samples sequenced in this study (Hadza and Nepali samples). Circles are samples from other studies. Distance matrix was generated with Jaccard similarity between samples. (**D**) Shannon diversity of H-G samples in our global metagenomics data set. Significance between groups was calculated using Wilcoxon rank-sum test (*** *P* < 0.001, * *P* < 0.05). (**E**) Confusion matrix from a random forest classifier built to predict the lifestyle of Hadza samples from this study and H-G and transitional lifestyle samples from publicly available studies. 100% of Hadza samples were classified as Hadza, H-G samples were correctly classified 53% of the time and transitional samples were classified correctly 91% of the time. (**F**) Scatter plot showing sequencing depth versus richness (number of observed species). Linear regression model of richness against sequencing depth reveals a highly significant association (*P* = 2.2 × 10^−16^). ‘H-G’, Hunter-Gatherer.

**Figure S5.**
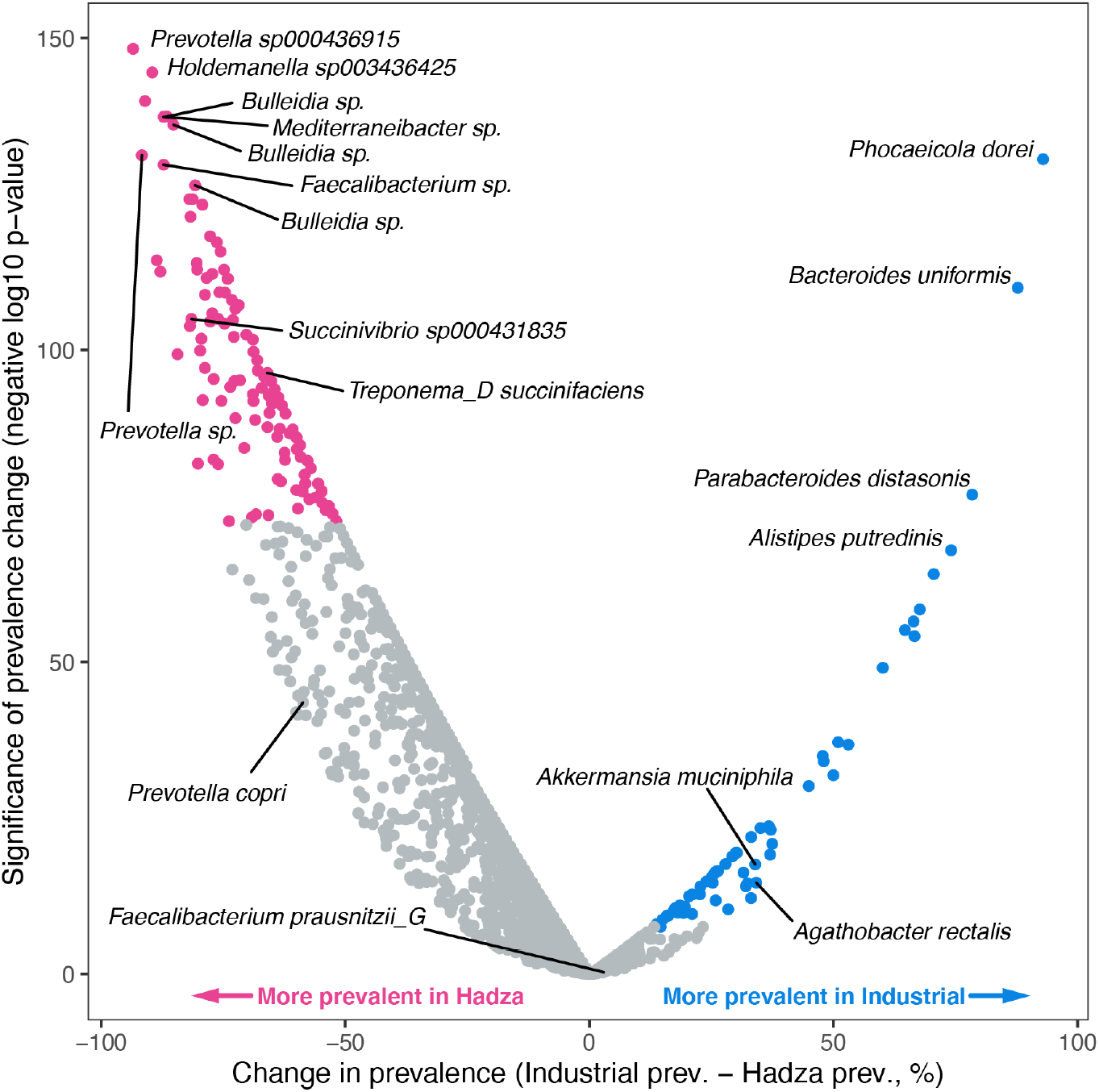
Lifestyle-specific enrichment of bacterial and archaeal taxa. Volcano plot showing enrichment of each species in either Hadza or industrial samples in our global metagenomics data set. Dots colored magenta are in the 95th percentile most enriched in the Hadza and are deemed VANISH taxa (124 total). Dots that are colored blue are in the 95th percentile most enriched in industrial samples and are deemed BloSSUM (63 total).

**Figure S6.**
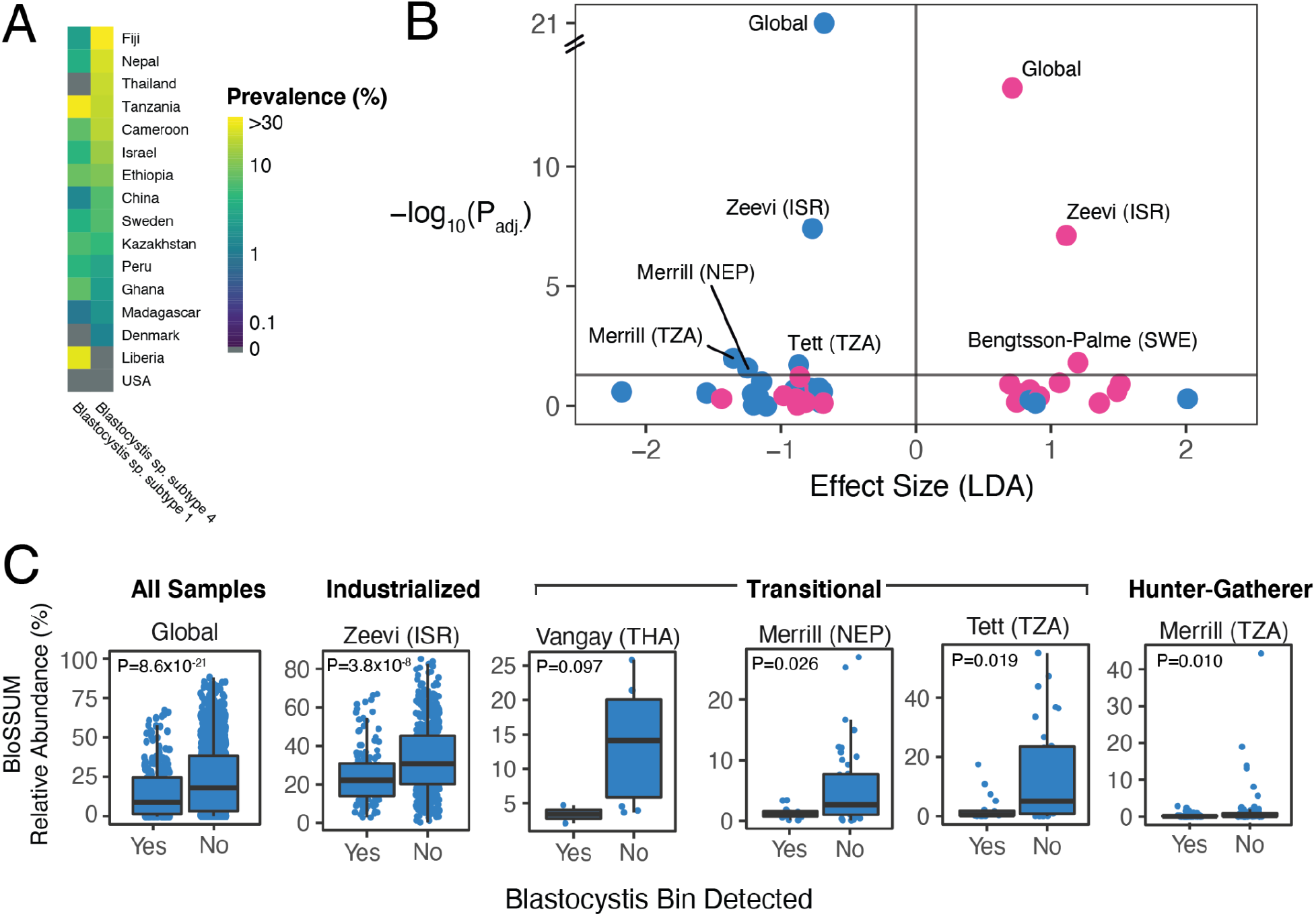
Global prevalence of two most prevalent *Blastocystis* MAGs and their association with VANISH and BloSSUM taxa. (**A**) A heatmap showing the prevalence of the two most prevalent *Blastocystis* MAGs (subtype 1 and subtype 4) in 16 different countries in our global metagenomics database. (**B**) A volcano plot showing associations between the presence or absence of either *Blastocystis* genome and the relative abundance of VANISH (magenta) or BloSSUM (blue) taxa. *P*-values were determined with a Wilcoxon rank-sum test and then adjusted with the Benjamini-Hochberg method to correct for multiple hypothesis testing. Threshold for significance of the adjusted *p-value*s is *P*=0.05 (or −log10(*P*)=1.3). Effect size was determined by linear discriminant analysis. The data points labeled “Global” are the associations for all samples in our global metagenomics data set. Other data points are for individual studies within the global metagenomics data set (annotated by first author of study and country of origin of the metagenomes). Across all studies we found that Blastocystis presence was positively and negatively associated with the total abundance of VANISH (*P*=5.1×10^−14^) and BloSSUM (*P*=8.6×10^−21^) taxa, respectively. (**C**) Boxplots showing the summed relative abundance of BloSSUM taxa per sample and whether *Blastocystis* was detected in that sample. Associations shown are for the entire global metagenomics data set and 5 additional populations (NEP = Nepal, TZA = Tanzania, THA = Thailand, ISR = Israel) from three lifestyles labeled above the plots. P-values shown are the results of Wilcoxon rank-sum tests.

**Figure S7.**
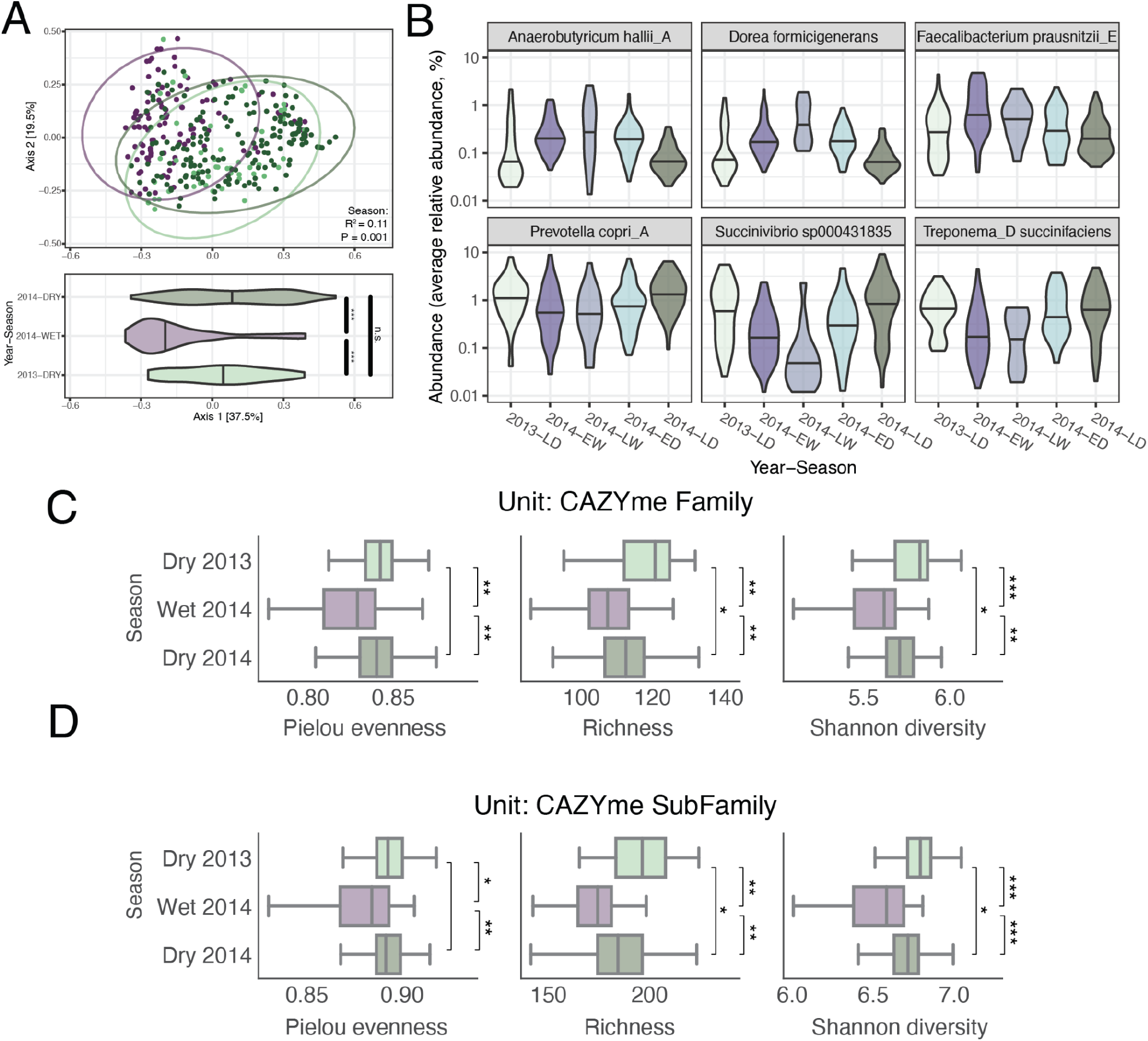
Seasonality in Hadza gut microbiome. **(A)** A principal coordinate analysis of Hadza samples where the Bray-Curtis distance matrix was generated by calculating the relative abundance of each taxonomic family in our bacterial/archaeal species-level genome database using InStrain (top panel). Samples are colored by season. Season explains a significant amount of the variation in the data (P = 0.001, R^2^ =0.09; ADONIS, using Subject ID as a strata). Sub-season also explains a significant amount of variation in the data (P = 0.001, R^2^ =0.14; ADONIS, using Subject ID as a strata). The bottom panel shows a violin plot of each sample’s PCo1 position, grouped by season. Samples collected in the dry season are significantly different from the wet season (P = 1.2 × 10^−10^ and P = 2 × 10^−16^ for 2013-DRY:2014-WET and 2014-WET:2014-DRY comparisons, respectively; Wilcoxon test). The samples collected in each dry season do not differ significantly from each other (P=0.34; Wilcoxon test). **(B)** The violin plots depict distribution of relative abundance for 6 SRGs that vary significantly over the 5 sub-seasons. The top three sub-panels depict species-level groups that have higher abundance in the wet seasons. The bottom three sub-panels depict species-level groups that have higher abundance in the dry seasons. (*Bulleidia sp*., P-adjusted = 7.5 × 10^−20^; *Dorea formicigenerans*, P-adjusted = 1.2 × 10^−16^; *Holdemanella sp003436425*, P-adjusted = 6.5 × 10^−16^; *Prevotella copri_*A, P-adjusted = 0.0054; *Succinivibrio sp000431835*, P-adjusted = 4.7 × 10^−7^; *Treponema_D succinifaciens*, P-adjusted = 0.012; Kruksal-Wallis test). 2013-LD (Late Dry); 2014-EW (Early Wet); 2014-LW (Late Wet); 2014-ED (Early Dry); 2014-LD (Late Dry). (**C**) For Hadza gut metagenomes sequenced in this study, genes present (≥ 80% breadth of coverage) on detected genomes (≥ 50% breadth of coverage) were annotated against the CAZyme database. CAZyme Pielou evenness (left), total richness (middle), and Shannon diversity (right) were calculated using the summed relative abundance of genomes containing each GH and PL CAZyme Family (for example, ‘GH16’) or (**D**) SubFamily (for example, ‘GH16.7’). Values for samples collected from Hadza individuals in different seasons were compared using a two-sided Wilcoxon rank-sum test (* P < 0.01; ** P < 1 × 10 ^−5^; *** P < 1 × 10^−10^).

## References

Abdill, R.J., Adamowicz, E.M., and Blekhman, R. (2022). Public human microbiome data are dominated by highly developed countries. PLoS Biol. 20, e3001536. .

Almeida, A., Mitchell, A.L., Boland, M., Forster, S.C., Gloor, G.B., Tarkowska, A., Lawley, T.D., and Finn, R.D. (2019). A new genomic blueprint of the human gut microbiota. Nature 568, 499–504. .

Almeida, A., Nayfach, S., Boland, M., Strozzi, F., Beracochea, M., Shi, Z.J., Pollard, K.S., Sakharova, E., Parks, D.H., Hugenholtz, P., et al. (2020). A unified catalog of 204,938 reference genomes from the human gut microbiome. Nat. Biotechnol. https://doi.org/10.1038/s41587-020-0603-3.

Andrews, S. (2010). FastQC: a quality control tool for high throughput sequence data.

Bäckhed, F., Roswall, J., Peng, Y., Feng, Q., Jia, H., Kovatcheva-Datchary, P., Li, Y., Xia, Y., Xie, H., Zhong, H., et al. (2015). Dynamics and Stabilization of the Human Gut Microbiome during the First Year of Life. Cell Host Microbe 17, 690–703. .

Belkhou, C., Tadeo, R.T., Bacigalupe, R., Valles-Colomer, M., Chaffron, S., Joossens, M., Obregon, A., Marín Reyes, L., Trujillo, O., Huys, G.R.B., et al. (2021). Treponema peruense sp. nov., a commensal spirochaete isolated from human faeces. Int. J. Syst. Evol. Microbiol. 71. https://doi.org/10.1099/ijsem.0.005050.

Bengtsson-Palme, J., Angelin, M., Huss, M., Kjellqvist, S., Kristiansson, E., Palmgren, H., Larsson, D.G.J., and Johansson, A. (2015). The Human Gut Microbiome as a Transporter of Antibiotic Resistance Genes between Continents. Antimicrob. Agents Chemother. 59, 6551–6560. .

Bland, C., Ramsey, T.L., Sabree, F., Lowe, M., Brown, K., Kyrpides, N.C., and Hugenholtz, P. (2007). CRISPR recognition tool (CRT): a tool for automatic detection of clustered regularly interspaced palindromic repeats. BMC Bioinformatics 8, 209. .

Blaser, M.J. (2017). The theory of disappearing microbiota and the epidemics of chronic diseases. Nat. Rev. Immunol. 17, 461–463. .

Blaser, M.J., and Falkow, S. (2009). What are the consequences of the disappearing human microbiota? Nat. Rev. Microbiol. 7, 887–894. .

Borges, R., Machado, J.P., Gomes, C., Rocha, A.P., and Antunes, A. (2019). Measuring phylogenetic signal between categorical traits and phylogenies. Bioinformatics 35, 1862–1869. .

Bowers, R.M., Kyrpides, N.C., Stepanauskas, R., Harmon-Smith, M., Doud, D., Reddy, T.B.K., Schulz, F., Jarett, J., Rivers, A.R., Eloe-Fadrosh, E.A., et al. (2017). Minimum information about a single amplified genome (MISAG) and a metagenome-assembled genome (MIMAG) of bacteria and archaea. Nat. Biotechnol. 35, 725–731. .

Brito, I.L., Yilmaz, S., Huang, K., Xu, L., Jupiter, S.D., Jenkins, A.P., Naisilisili, W., Tamminen, M., Smillie, C.S., Wortman, J.R., et al. (2016). Mobile genes in the human microbiome are structured from global to individual scales. Nature https://doi.org/10.1038/nature18927.

Brown, C.T., Olm, M.R., Thomas, B.C., and Banfield, J.F. (2016). Measurement of bacterial replication rates in microbial communities. Nat. Biotechnol. 34, 1256–1263. .

Buchfink, B., Xie, C., and Huson, D.H. (2015). Fast and sensitive protein alignment using DIAMOND. Nat. Methods 12, 59–60. .

Bushnell, B. (2014). BBTools software package. URL Http://sourceforge.Net/projects/bbmap 578, 579. .

Bushnell, B., Rood, J., and Singer, E. (2017). BBMerge--accurate paired shotgun read merging via overlap. PLoS One 12, e0185056. .

Camacho, C., Coulouris, G., Avagyan, V., Ma, N., Papadopoulos, J., Bealer, K., and Madden, T.L. (2009). BLAST+: architecture and applications. BMC Bioinformatics 10, 421. .

Chaumeil, P.-A., Mussig, A.J., Hugenholtz, P., and Parks, D.H. (2019). GTDB-Tk: a toolkit to classify genomes with the Genome Taxonomy Database. Bioinformatics https://doi.org/10.1093/bioinformatics/btz848.

Chen, Y., Ye, W., Zhang, Y., and Xu, Y. (2015). High speed BLASTN: an accelerated MegaBLAST search tool. Nucleic Acids Res. 43, 7762–7768. .

Clark, C.G., van der Giezen, M., Alfellani, M.A., and Stensvold, C.R. (2013). Recent developments in Blastocystis research. Adv. Parasitol. 82, 1–32. .

Clemente, J.C., Pehrsson, E.C., Blaser, M.J., Sandhu, K., Gao, Z., Wang, B., Magris, M., Hidalgo, G., Contreras, M., Noya-Alarcón, Ó., et al. (2015). The microbiome of uncontacted Amerindians. Sci Adv 1. https://doi.org/10.1126/sciadv.1500183.

Conteville, L.C., Oliveira-Ferreira, J., and Vicente, A.C.P. (2019). Gut Microbiome Biomarkers and Functional Diversity Within an Amazonian Semi-Nomadic Hunter-Gatherer Group. Front. Microbiol. 10. https://doi.org/10.3389/fmicb.2019.01743.

Costea, P.I., Coelho, L.P., Sunagawa, S., Munch, R., Huerta-Cepas, J., Forslund, K., Hildebrand, F., Kushugulova, A., Zeller, G., and Bork, P. (2017a). Subspecies in the global human gut microbiome. Mol. Syst. Biol. 13, 960. .

Costea, P.I., Coelho, L.P., Sunagawa, S., Munch, R., Huerta-Cepas, J., Forslund, K., Hildebrand, F., Kushugulova, A., Zeller, G., and Bork, P. (2017b). Subspecies in the global human gut microbiome. Mol. Syst. Biol. 13, 960. .

Dixon, P. (2003). VEGAN, a package of R functions for community ecology. J. Veg. Sci. 14, 927–930. .

Eddy, S.R. (2011). Accelerated Profile HMM Searches. PLoS Comput. Biol. 7, e1002195. .

Edgar, R.C. (2007). PILER-CR: fast and accurate identification of CRISPR repeats. BMC Bioinformatics 8, 18. .

Edgar, R.C. (2021). MUSCLE v5 enables improved estimates of phylogenetic tree confidence by ensemble bootstrapping.

El-Gebali, S., Mistry, J., Bateman, A., Eddy, S.R., Luciani, A., Potter, S.C., Qureshi, M., Richardson, L.J., Salazar, G.A., Smart, A., et al. (2018). The Pfam protein families database in 2019. Nucleic Acids Res. https://doi.org/10.1093/nar/gky995.

Eren, A.M., Esen, Ö.C., Quince, C., Vineis, J.H., Morrison, H.G., Sogin, M.L., and Delmont, T.O. (2015). Anvi’o: an advanced analysis and visualization platform for ‘omics data. PeerJ 3, e1319. .

Faith, J.J., Guruge, J.L., Charbonneau, M., Subramanian, S., Seedorf, H., Goodman, A.L., Clemente, J.C., Knight, R., Heath, A.C., Leibel, R.L., et al. (2013). The Long-Term Stability of the Human Gut Microbiota. Science 341, 1237439–1237439. .

Falush, D., Wirth, T., Linz, B., Pritchard, J.K., Stephens, M., Kidd, M., Blaser, M.J., Graham, D.Y., Vacher, S., Perez-Perez, G.I., et al. (2003). Traces of human migrations in Helicobacter pylori populations. Science 299, 1582–1585. .

Fragiadakis, G.K., Smits, S.A., Sonnenburg, E.D., Van Treuren, W., Reid, G., Knight, R., Manjurano, A., Changalucha, J., Dominguez-Bello, M.G., Leach, J., et al. (2019). Links between environment, diet, and the hunter-gatherer microbiome. Gut Microbes 10, 216–227. .

Green, E.D., Gunter, C., Biesecker, L.G., Di Francesco, V., Easter, C.L., Feingold, E.A., Felsenfeld, A.L., Kaufman, D.J., Ostrander, E.A., Pavan, W.J., et al. (2020). Strategic vision for improving human health at The Forefront of Genomics. Nature 586, 683–692. .

Groussin, M., Poyet, M., Sistiaga, A., Kearney, S.M., Moniz, K., Noel, M., Hooker, J., Gibbons, S.M., Segurel, L., Froment, A., et al. (2021). Elevated rates of horizontal gene transfer in the industrialized human microbiome. Cell 184, 2053–2067.e18. .

Gurevich, A., Saveliev, V., Vyahhi, N., and Tesler, G. (2013). QUAST: quality assessment tool for genome assemblies. Bioinformatics 29, 1072–1075. .

Gutierrez, S.C., Martinez, J.M.S., and Gabaldón, T. (2009). TrimAl: a Tool for automatic alignment trimming. Bioinformatics 25, 1972–1973. .

Hockenberry, A.J., and Wilke, C.O. (2021). BACPHLIP: predicting bacteriophage lifestyle from conserved protein domains. PeerJ 9, e11396. .

Human Microbiome Project Consortium (2012). Structure, function and diversity of the healthy human microbiome. Nature 486, 207–214. .

Hyatt, D., Chen, G.-L., LoCascio, P.F., Land, M.L., Larimer, F.W., and Hauser, L.J. (2010). Prodigal: prokaryotic gene recognition and translation initiation site identification. BMC Bioinformatics 11, 119. .

Jha, A.R., Davenport, E.R., Gautam, Y., Bhandari, D., Tandukar, S., Ng, K.M., Fragiadakis, G.K., Holmes, S., Gautam, G.P., Leach, J., et al. (2018). Gut microbiome transition across a lifestyle gradient in Himalaya. PLoS Biol. 16, e2005396. .

Jones, E., Oliphant, T., and Peterson, P. (2001). SciPy: Open source scientific tools for Python. URL Http://scipy.Org.

Kang, D.D., Li, F., Kirton, E., Thomas, A., Egan, R., An, H., and Wang, Z. (2019). MetaBAT 2: an adaptive binning algorithm for robust and efficient genome reconstruction from metagenome assemblies. PeerJ 7, e7359. .

Langmead, B., and Salzberg, S.L. (2012). Fast gapped-read alignment with Bowtie 2. Nat. Methods 9, 357–359. .

Lee, M.D. (2019). GToTree: a user-friendly workflow for phylogenomics. Bioinformatics 35, 4162–4164. .

Letunic, I., and Bork, P. (2007). Interactive Tree Of Life (iTOL): an online tool for phylogenetic tree display and annotation. Bioinformatics 23, 127–128. .

Lewis, T.E., Sillitoe, I., and Lees, J.G. (2019). cath-resolve-hits: a new tool that resolves domain matches suspiciously quickly. Bioinformatics 35, 1766–1767. .

Lind, A.L., and Pollard, K.S. (2021). Accurate and sensitive detection of microbial eukaryotes from whole metagenome shotgun sequencing. Microbiome 9, 58. .

Linz, B., Balloux, F., Moodley, Y., Manica, A., Liu, H., Roumagnac, P., Falush, D., Stamer, C., Prugnolle, F., van der Merwe, S.W., et al. (2007). An African origin for the intimate association between humans and Helicobacter pylori. Nature 445, 915–918. .

Litvak, Y., Byndloss, M.X., and Bäumler, A.J. (2018). Colonocyte metabolism shapes the gut microbiota. Science 362. https://doi.org/10.1126/science.aat9076.

Liu, H., Prugnolle, F., Manica, A., and Balloux, F. (2006). A geographically explicit genetic model of worldwide human-settlement history. Am. J. Hum. Genet. 79, 230–237. .

Liu, W., Zhang, J., Wu, C., Cai, S., Huang, W., Chen, J., Xi, X., Liang, Z., Hou, Q., Zhou, B., et al. (2016). Unique Features of Ethnic Mongolian Gut Microbiome revealed by metagenomic analysis. Sci. Rep. 6, 34826. .

Liu, W., Zhang, J., Wu, C., Cai, S., Huang, W., Chen, J., Xi, X., Liang, Z., Hou, Q., Zhou, B., et al. (2017). Corrigendum: Unique Features of Ethnic Mongolian Gut Microbiome revealed by metagenomic analysis. Sci. Rep. 7, 39576. .

Lizano, S., Luo, F., and Bessen, D.E. (2007). Role of streptococcal T antigens in superficial skin infection. J. Bacteriol. 189, 1426–1434. .

Lloyd-Price, J., Mahurkar, A., Rahnavard, G., Crabtree, J., Orvis, J., Hall, A.B., Brady, A., Creasy, H.H., McCracken, C., Giglio, M.G., et al. (2017). Strains, functions and dynamics in the expanded Human Microbiome Project. Nature https://doi.org/10.1038/nature23889.

Lokmer, A., Cian, A., Froment, A., Gantois, N., Viscogliosi, E., Chabé, M., and Ségurel, L. (2019). Use of shotgun metagenomics for the identification of protozoa in the gut microbiota of healthy individuals from worldwide populations with various industrialization levels. PLoS One 14, e0211139. .

Mangola, S.M., Lund, J.R., Schnorr, S.L., and Crittenden, A.N. (2022). Ethical microbiome research with Indigenous communities. Nat Microbiol 7, 749–756. .

Marlowe, F. (2010). The Hadza: Hunter-gatherers of Tanzania (University of California Press).

Marlowe, F.W., Berbesque, J.C., Wood, B., Crittenden, A., Porter, C., and Mabulla, A. (2014). Honey, Hadza, hunter-gatherers, and human evolution. J. Hum. Evol. 71, 119–128. .

Martínez, I., Stegen, J.C., Maldonado-Gómez, M.X., Eren, A.M., Siba, P.M., Greenhill, A.R., and Walter, J. (2015). The gut microbiota of rural papua new guineans: composition, diversity patterns, and ecological processes. Cell Rep. 11, 527–538. .

Minh, B.Q., Schmidt, H.A., Chernomor, O., Schrempf, D., Woodhams, M.D., von Haeseler, A., and Lanfear, R. (2020). IQ-TREE 2: New Models and Efficient Methods for Phylogenetic Inference in the Genomic Era. Mol. Biol. Evol. 37, 1530–1534. .

Mistry, J., Chuguransky, S., Williams, L., Qureshi, M., Salazar, G.A., Sonnhammer, E.L.L., Tosatto, S.C.E., Paladin, L., Raj, S., Richardson, L.J., et al. (2021). Pfam: The protein families database in 2021. Nucleic Acids Res. 49, D412–D419. .

Modi, S.R., Collins, J.J., and Relman, D.A. (2014). Antibiotics and the gut microbiota. J. Clin. Invest. 124, 4212–4218. .

Moeller, A.H., Li, Y., Mpoudi Ngole, E., Ahuka-Mundeke, S., Lonsdorf, E.V., Pusey, A.E., Peeters, M., Hahn, B.H., and Ochman, H. (2014). Rapid changes in the gut microbiome during human evolution. Proc. Natl. Acad. Sci. U. S. A. 111, 16431–16435. .

Mueller, N.T., Bakacs, E., Combellick, J., Grigoryan, Z., and Dominguez-Bello, M.G. (2015). The infant microbiome development: mom matters. Trends Mol. Med. 21, 109–117. .

Nayfach, S., Shi, Z.J., Seshadri, R., Pollard, K.S., and Kyrpides, N.C. (2019). New insights from uncultivated genomes of the global human gut microbiome. Nature https://doi.org/10.1038/s41586-019-1058-x.

Nayfach, S., Camargo, A.P., Schulz, F., Eloe-Fadrosh, E., Roux, S., and Kyrpides, N.C. (2021a). CheckV assesses the quality and completeness of metagenome-assembled viral genomes. Nat. Biotechnol. 39, 578–585. .

Nayfach, S., Páez-Espino, D., Call, L., Low, S.J., Sberro, H., Ivanova, N.N., Proal, A.D., Fischbach, M.A., Bhatt, A.S., Hugenholtz, P., et al. (2021b). Metagenomic compendium of 189,680 DNA viruses from the human gut microbiome. Nat Microbiol 6, 960–970. .

NCBI Resource Coordinators (2017). Database Resources of the National Center for Biotechnology Information. Nucleic Acids Res. 45, D12–D17. .

Nurk, S., Meleshko, D., Korobeynikov, A., and Pevzner, P.A. (2017). metaSPAdes: a new versatile metagenomic assembler. Genome Res. 27, 824–834. .

Obregon-Tito, A.J., Tito, R.Y., Metcalf, J., Sankaranarayanan, K., Clemente, J.C., Ursell, L.K., Zech Xu, Z., Van Treuren, W., Knight, R., Gaffney, P.M., et al. (2015). Subsistence strategies in traditional societies distinguish gut microbiomes. Nat. Commun. 6, 6505. .

Olm, M.R., Brown, C.T., Brooks, B., and Banfield, J.F. (2017). dRep: a tool for fast and accurate genomic comparisons that enables improved genome recovery from metagenomes through de-replication. ISME J. 11, 2864–2868. .

Olm, M.R., West, P.T., Brooks, B., Firek, B.A., Baker, R., Morowitz, M.J., and Banfield, J.F. (2019a). Genome-resolved metagenomics of eukaryotic populations during early colonization of premature infants and in hospital rooms. Microbiome 7, 26. .

Olm, M.R., Bhattacharya, N., Crits-Christoph, A., Firek, B.A., Baker, R., Song, Y.S., Morowitz, M.J., and Banfield, J.F. (2019b). Necrotizing enterocolitis is preceded by increased gut bacterial replication, Klebsiella, and fimbriae-encoding bacteria. Science Advances 5, eaax5727. .

Olm, M.R., Crits-Christoph, A., Diamond, S., Lavy, A., Matheus Carnevali, P.B., and Banfield, J.F. (2020). Consistent Metagenome-Derived Metrics Verify and Delineate Bacterial Species Boundaries. mSystems 5. https://doi.org/10.1128/mSystems.00731-19.

Olm, M.R., Crits-Christoph, A., Bouma-Gregson, K., Firek, B.A., Morowitz, M.J., and Banfield, J.F. (2021). inStrain profiles population microdiversity from metagenomic data and sensitively detects shared microbial strains. Nat. Biotechnol. https://doi.org/10.1038/s41587-020-00797-0.

Olm, M.R., Dahan, D., Carter, M.M., Merrill, B.D., Yu, F.B., Jain, S., Meng, X., Tripathi, S., Wastyk, H., Neff, N., et al. (2022). Robust variation in infant gut microbiome assembly across a spectrum of lifestyles. Science 376, 1220–1223. .

Ondov, B.D., Treangen, T.J., Melsted, P., Mallonee, A.B., Bergman, N.H., Koren, S., and Phillippy, A.M. (2016). Mash: fast genome and metagenome distance estimation using MinHash. Genome Biol. 17. https://doi.org/10.1186/s13059-016-0997-x.

Parks, D.H., Imelfort, M., Skennerton, C.T., Hugenholtz, P., and Tyson, G.W. (2015). CheckM: assessing the quality of microbial genomes recovered from isolates, single cells, and metagenomes. Genome Res. 25, 1043–1055. .

Parks, D.H., Chuvochina, M., Rinke, C., Mussig, A.J., Chaumeil, P.-A., and Hugenholtz, P. (2022). GTDB: an ongoing census of bacterial and archaeal diversity through a phylogenetically consistent, rank normalized and complete genome-based taxonomy. Nucleic Acids Res. 50, D785–D794. .

Pasolli, E., Asnicar, F., Manara, S., Zolfo, M., Karcher, N., Armanini, F., Beghini, F., Manghi, P., Tett, A., Ghensi, P., et al. (2019). Extensive Unexplored Human Microbiome Diversity Revealed by Over 150,000 Genomes from Metagenomes Spanning Age, Geography, and Lifestyle. Cell 176, 649–662.e20. .

Pehrsson, E.C., Tsukayama, P., Patel, S., Mejía-Bautista, M., Sosa-Soto, G., Navarrete, K.M., Calderon, M., Cabrera, L., Hoyos-Arango, W., Bertoli, M.T., et al. (2016). Interconnected microbiomes and resistomes in low-income human habitats. Nature 533, 212–216. .

Prjibelski, A., Antipov, D., Meleshko, D., Lapidus, A., and Korobeynikov, A. (2020). Using SPAdes DE Novo Assembler. Curr. Protoc. Bioinformatics 70, e102. .

Qin, J., Li, R., Raes, J., Arumugam, M., Burgdorf, K.S., Manichanh, C., Nielsen, T., Pons, N., Levenez, F., Yamada, T., et al. (2010). A human gut microbial gene catalogue established by metagenomic sequencing. Nature 464, 59–65. .

Qin, J., Li, Y., Cai, Z., Li, S., Zhu, J., Zhang, F., Liang, S., Zhang, W., Guan, Y., Shen, D., et al. (2012). A metagenome-wide association study of gut microbiota in type 2 diabetes. Nature 490, 55–60. .

Rampelli, S., Schnorr, S.L., Consolandi, C., Turroni, S., Severgnini, M., Peano, C., Brigidi, P., Crittenden, A.N., Henry, A.G., and Candela, M. (2015). Metagenome Sequencing of the Hadza Hunter-Gatherer Gut Microbiota. Curr. Biol. 25, 1682–1693. .

Revell, L.J. (2012). phytools: an R package for phylogenetic comparative biology (and other things). Methods Ecol. Evol. 3, 217–223. .

Rodriguez-Valera, F., Martin-Cuadrado, A.-B., Rodriguez-Brito, B., Pašić, L., Thingstad, T.F., Rohwer, F., and Mira, A. (2009). Explaining microbial population genomics through phage predation. Nat. Rev. Microbiol. 7, 828–836. .

Rosa, B.A., Supali, T., Gankpala, L., Djuardi, Y., Sartono, E., Zhou, Y., Fischer, K., Martin, J., Tyagi, R., Bolay, F.K., et al. (2018). Differential human gut microbiome assemblages during soil-transmitted helminth infections in Indonesia and Liberia. Microbiome 6, 33. .

Saary, P., Mitchell, A.L., and Finn, R.D. (2020). Estimating the quality of eukaryotic genomes recovered from metagenomic analysis with EukCC. Genome Biol. 21, 244. .

Seabold, S., and Perktold, J. (2010). Statsmodels: Econometric and statistical modeling with python. In Proceedings of the 9th Python in Science Conference, (Austin, TX), p. 61.

Smits, S.A., Leach, J., Sonnenburg, E.D., Gonzalez, C.G., Lichtman, J.S., Reid, G., Knight, R., Manjurano, A., Changalucha, J., Elias, J.E., et al. (2017). Seasonal cycling in the gut microbiome of the Hadza hunter-gatherers of Tanzania. Science 357, 802–806. .

Sonnenburg, E.D., and Sonnenburg, J.L. (2019a). The ancestral and industrialized gut microbiota and implications for human health. Nat. Rev. Microbiol. 17, 383–390. .

Sonnenburg, J.L., and Sonnenburg, E.D. (2019b). Vulnerability of the industrialized microbiota. Science 366. https://doi.org/10.1126/science.aaw9255.

Steinegger, M., and Söding, J. (2017). MMseqs2 enables sensitive protein sequence searching for the analysis of massive data sets. Nat. Biotechnol. 35, 1026–1028. .

Suzek, B.E., Huang, H., McGarvey, P., Mazumder, R., and Wu, C.H. (2007). UniRef: comprehensive and non-redundant UniProt reference clusters. Bioinformatics 23, 1282–1288. .

Suzuki, T.A., Fitzstevens, L., Schmidt, V.T., Enav, H., Huus, K., Mbong, M., Adegbite, B.R., Zinsou, J.F., Esen, M., Velavan, T.P., et al. (2021). Codiversification of gut microbiota with humans.

Tamburini, F.B., Maghini, D., Oduaran, O.H., Brewster, R., Hulley, M.R., Sahibdeen, V., Norris, S.A., Tollman, S., Kahn, K., Wagner, R.G., et al. (2022). Short- and long-read metagenomics of urban and rural South African gut microbiomes reveal a transitional composition and undescribed taxa. Nat. Commun. 13, 926. .

Tange, O. (2018). GNU Parallel 2018 (Lulu.com).

Tett, A., Huang, K.D., Asnicar, F., Fehlner-Peach, H., Pasolli, E., Karcher, N., Armanini, F., Manghi, P., Bonham, K., Zolfo, M., et al. (2019). The Prevotella copri Complex Comprises Four Distinct Clades Underrepresented in Westernized Populations. Cell Host Microbe 26, 666–679.e7. .

Valles-Colomer, M., Bacigalupe, R., Vieira-Silva, S., Suzuki, S., Darzi, Y., Tito, R.Y., Yamada, T., Segata, N., Raes, J., and Falony, G. (2022). Variation and transmission of the human gut microbiota across multiple familial generations. Nat Microbiol 7, 87–96. .

Vangay, P., Johnson, A.J., Ward, T.L., Al-Ghalith, G.A., Shields-Cutler, R.R., Hillmann, B.M., Lucas, S.K., Beura, L.K., Thompson, E.A., Till, L.M., et al. (2018). US Immigration Westernizes the Human Gut Microbiome. Cell 175, 962–972.e10. .

Waskom, M. (2021). seaborn: statistical data visualization. J. Open Source Softw. 6, 3021. .

Wastyk, H.C., Fragiadakis, G.K., Perelman, D., Dahan, D., Merrill, B.D., Yu, F.B., Topf, M., Gonzalez, C.G., Van Treuren, W., Han, S., et al. (2021). Gut-microbiota-targeted diets modulate human immune status. Cell 184, 4137–4153.e14. .

West, P.T., Probst, A.J., Grigoriev, I.V., Thomas, B.C., and Banfield, J.F. (2018). Genome-reconstruction for eukaryotes from complex natural microbial communities. Genome Res. 28, 569–580. .

Wibowo, M.C., Yang, Z., Borry, M., Hübner, A., Huang, K.D., Tierney, B.T., Zimmerman, S., Barajas-Olmos, F., Contreras-Cubas, C., García-Ortiz, H., et al. (2021). Reconstruction of ancient microbial genomes from the human gut. Nature 594, 234–239. .

Yatsunenko, T., Rey, F.E., Manary, M.J., Trehan, I., Dominguez-Bello, M.G., Contreras, M., Magris, M., Hidalgo, G., Baldassano, R.N., Anokhin, A.P., et al. (2012). Human gut microbiome viewed across age and geography. Nature https://doi.org/10.1038/nature11053.

Zeevi, D., Korem, T., Zmora, N., Israeli, D., Rothschild, D., Weinberger, A., Ben-Yacov, O., Lador, D., Avnit-Sagi, T., Lotan-Pompan, M., et al. (2015). Personalized Nutrition by Prediction of Glycemic Responses. Cell 163, 1079–1094. .

Zhang, H., Yohe, T., Huang, L., Entwistle, S., Wu, P., Yang, Z., Busk, P.K., Xu, Y., and Yin, Y. (2018). dbCAN2: a meta server for automated carbohydrate-active enzyme annotation. Nucleic Acids Res. 46, W95–W101. .

